# Context-Aware Hydrophobicity Modeling: HydroMap and FastHydroMap

**DOI:** 10.64898/2026.06.07.730647

**Authors:** Samuel Lobo, Saeed Najafi, Joan-Emma Shea, M. Scott Shell

## Abstract

Hydrophobicity governs a vast range of phenomena, from protein–protein interactions to nanomaterial assembly, and can be rigorously quantified by the dewetting free energy (Fdewet) of a molecule or surface. However, hydrophobicity remains widely treated as an additive property of amino acid identity, obscuring the fact that water’s response is a collective property of the surface, shaped by curvature, chemical patterning, and neighboring residues. Direct calculation of Fdewet via specialized molecular simulations captures this collective behavior but is prohibitively slow, leaving in place broadly-used, decades-old sequence-based hydropathy scales that neglect the physics of solvation. Here we show that residue-level Fdewet can be predicted from a compact set of local water features (water structural signatures and residue–water potential energy) extracted from a brief and inexpensive all-atom simulation. We embed this insight in two models: HydroMap, which predicts Fdewet directly from water features, and FastHydroMap, a computationally inexpensive graph neural network surrogate trained on HydroMap that requires no solvent simulation. HydroMap and FastHydroMap capture context-dependent hydrophobicity that classical, sequence-only hydropathy scales miss. We demonstrate this across three protein systems: on an α-synuclein amyloid filament, strongly dewetting interfaces align with unassigned peptide densities, revealing hidden binding sites; in calmodulin, hydrophobicity redistributes upon Ca^2+^ binding; and for Protein G, time-resolved hydrophobicity changes track the folding trajectory. Together, these models make Fdewet a computationally inexpensive descriptor for proteins, membranes, and other surfaces, enabling rapid scoring for materials design and a time-resolved view of dynamic hydrophobic-mediated processes such as protein folding.

**Significance Statement:** Hydrophobicity, the tendency of surfaces to expel water, drives how proteins fold and how molecules recognize one another. For decades, it has widely been treated as a fixed property of an amino acid or chemical group, but water actually responds to the collective shape and chemistry of a surface, such as that presented by a protein, not to its components in isolation. Measuring this collective response from molecular simulation is rigorous but prohibitively slow. We show that it can instead be inferred from a compact set of features describing water structure and interactions near a surface, and we use this insight to build models that predict hydrophobicity rapidly and at residue resolution, enabling practical, physically grounded design of hydrophobic-mediated interactions.

## Introduction

Water is far from a passive medium in biology and materials science – it actively shapes molecular interactions through its collective structure and fluctuations. In aqueous environments, water-mediated hydrophobic interactions can drive proteins to fold and aggregate (1–4), viruses to assemble their capsids (5, 6), and polymers or colloids to cluster or phase separate (7–9). These processes, though spanning vastly different length scales, all emerge from the same underlying thermodynamic drive: water tends to minimize its contact with exposed hydrophobic surfaces. When two solvated solutes or surfaces approach one another, the water between them can be expelled in a process called “desolvating” or “dewetting”. This expulsion of interfacial water can be thermodynamically favorable, especially around nonpolar surfaces, thus stabilizing the interaction between the surfaces. By better understanding and quantifying the thermodynamic driving force of this dewetting process, we can better predict—and ultimately control—water-mediated association and assembly across biological and materials systems.

One particularly useful metric for the desolvating/dewetting process is the dewetting free energy (*F*_*dewet*_), which quantifies the thermodynamic cost of removing water from a particular spatially-defined region (such as a protein pocket or a material surface). Regions with a low (i.e. favorable) *F*_*dewet*_ are more prone to “dry out,” indicating stronger hydrophobic character, while regions with a high *F*_*dewet*_ can more tenaciously hold onto water, indicating hydrophilic character. Computing *F*_*dewet*_directly, however, is nontrivial as it requires evaluating the free energy change associated with removing water from a defined volume, essentially sampling the rare fluctuations in which that volume empties of water. Advanced molecular simulation techniques have been developed to accomplish such calculations. One prominent approach is Indirect Umbrella Sampling (INDUS), which biases the number of water molecules in a selected region to measure the free-energy cost of desolvating it (10–13). INDUS provides a physically rigorous way to map out hydrophobic hot spots on molecular surfaces by gradually desolvating waters from and observing interfaces that desolvate (13, 14), or by calculating the work required to desolvate specific interfaces (11). This approach has yielded valuable insight into hydrophobic interfaces – for example, revealing non-additive effects in chemically patterned surfaces (15), revealing hydrophobic patches relevant in protein-protein interactions (14), and allowing the definition of a dewetting-based hydrophobicity scale for amino acids that are influenced by its context (i.e. the neighboring amino acids and the local geometry) (3, 16). It is important to note that other approaches, notably topography-aware hydropathy scales, have also demonstrated that a residue’s hydropathic character depends strongly on local protein geometry (17), independently reinforcing the context-dependence of hydrophobicity.

In practice, coloring protein surfaces by hydrophobicity is already one of the most routine steps in structural biology — visualization of a protein’s 3D structure and painting it with a hydrophobicity scale is a common workflow to identify potential binding sites, aggregation-prone regions, or druggable pockets. Yet the conventional scales underlying these maps (e.g. Kyte-Doolittle(18)) assign a fixed value to each residue regardless of their structural and chemical environment: a phenylalanine in a concave pocket, on a convex exposed loop, or flanked by charged residues all receive the same score, yet such contexts produce measurably different dewetting propensities (19). Other approaches like the molecular lipophilicity potential (20, 21) and spatial aggregation propensity (22) go further by spatially averaging hydrophobic contributions within a local neighborhood, thereby capturing clustering of hydrophobic residues. However, these methods remain fundamentally additive – they sum fixed atomic or residue-level contributions and cannot capture the collective, non-additive reorganization that governs true surface hydrophobicity (19, 23). That is, hydrophobicity is a collective surface property, not an amino acid property.

INDUS F_dewet_ hydrophobicity maps would be far more informative than conventional scales, but generating them routinely remains impractical. INDUS must extensively sample water-density fluctuations, often needing dozens of simulations and hours to converge the free energy calculations. A separate set of simulations is typically required for each new interface—for example, each amino acid on a protein or each new mutation at a designed interface. This computational expense makes it difficult to screen large systems or many variants. For instance, mapping F_dewet_ at each residue of a large protein complex or scanning hundreds of possible surface chemistries would be impractical with INDUS. This practical gap strongly motivates faster surrogate methods to estimate dewetting/desolvating free energies with significantly less computational effort.

More fundamentally, there has been major interest in understanding the extent to which thermodynamic measures of hydrophobicity like F_dewet_ are signaled from underlying hydration water structure and interactions. Recent efforts have characterized a range of water order parameters that potentially encode dewetting propensity without running full free energy calculations, including water angle correlations. In particular, the water triplet distribution (also called the water 3-body angle distribution) and the tetrahedral order parameter *q* have been shown to correlate with a number of thermodynamic and kinetic properties of water (3, 16, 24–29). These metrics can capture shifts among tetrahedral (∼109.5°), ideal-gas-like (∼90°), and icosahedral (∼60°) motifs at interfaces, and the 48° water triplet feature has been linked to hyper-coordinated water and higher dewetting free energies (more hydrophilic) on patterned surfaces (30). Complementary structural descriptors probe the hydrogen-bond (H-bond) network, including water-water & water-solute H-bond statistics, and water ring statistics (e.g., tetra-, penta-, and hexameric rings) formed by H-bonded waters (31, 32). Translational descriptors include the local structure index (LSI), the fifth-neighbor distance d_5_, and the ζ order parameter, which gauge first- and second-shell intermolecular separation distances. Importantly, density/occupancy fluctuations P(N) within a defined shell provide a collective coordinate for wet–dry transitions (33–35): around extended hydrophobes the low-N tail is enhanced, while small hydrophobes retain near-bulk shell structure with only modest angular or H-bond reorganizations (36). A complementary geometric view connects cavity shapes (via Voronoi/alpha-shape analyses) to solvation thermodynamics, showing that void topology around amino-acid–like solutes tracks thermodynamic trends (37, 38).

Beyond simulation-computed water order parameters, data-driven surrogates have become powerful tools for predicting hydrophobicity. Convolutional neural networks trained on interfacial water configurations of chemically patterned SAMs can predict hydration free energies across large libraries of surface chemistries, revealing pronounced non-additive effects of polar– nonpolar arrangement that additive models miss (39–41). Taken together, these efforts motivate the idea of predicting local hydrophobicity with compact, interpretable features and ML surrogates to retain explicit-water insight while drastically reducing sampling.

Building on these developments, here we introduce HydroMap and FastHydroMap, fast and ultrafast surrogate models for predicting residue-level *F*_*dewet*_ for protein-based systems as a quantitative measure of local hydrophobicity. Guided by a simple solvation-thermodynamics picture, HydroMap builds on linear models with three features: the ensemble-averaged residue– water potential energy and two LASSO-selected water-structure fingerprints. HydroMap’s training data consists of amino acid INDUS F_dewet_ data in two contexts, isolated and protein-embedded, spanning a variety of chemistries and geometric environments. We additionally compute F_dewet_ for isolated amino acids across three force fields where *F*_dewet_ varies significantly due to differences in water structuring and protein–water interactions (42). After training, HydroMap predictions then involve a short, unbiased all-atom MD simulation from which we extract residue–water potential energies and local water structural features. This yields an interpretable, fast model that recovers residue-level INDUS F_dewet_ across contexts and provides the ability to generate larger labelled datasets for training further, ultrafast prediction algorithms.

Having established an accurate, interpretable model, we then significantly accelerate hydrophobicity calculation by training FastHydroMap on HydroMap outputs to predict F_dewet_ with a graph neural network, without an MD simulation. This accelerates F_dewet_ calculations by approximately three orders-of-magnitude, enabling high throughput analyses. FastHydroMap is a message passing neural network trained from tens of thousands of amino acid HydroMap *F*_*dewet*_ predictions across nearly 1000 small, structured proteins. Overall, the two F_dewet_ predictors in this work are several orders of magnitude faster than traditional INDUS-based calculations – fast enough to evaluate *F*_*dewet*_ for every residue in a large protein in tens-to-hundreds of seconds (HydroMap) or fractions of a second (FastHydroMap). This enables these methods to screen many thousands of sites across different conditions and quantify context-aware hydrophobicity.

Collectively, HydroMap and FastHydroMap provide computationally efficient, context-aware estimates of a surface’s propensity to desolvate – capturing the dependence of each residue’s hydrophobicity on its local geometric and chemical environment that conventional scales neglect. We showcase their applications in a variety of systems with static proteins, a conformationally plastic fold-switching protein, and a dynamic protein folding trajectory. Specifically, we measure hydrophobicity in (a) the Parkinson’s amyloid filament, (b) the fold-switching calmodulin protein, and (c) Protein G’s dynamic folding trajectory. Our identification of three minimal water features—one energetic and two structural water descriptors—reinforces a fundamental link between water structure and hydrophobicity thermodynamics. Because FastHydroMap evaluates an entire protein surface in a fraction of a second at negligible computational cost, it is a practical candidate to replace conventional hydropathy-scale coloring of protein surfaces with physics-informed, context-dependent hydrophobicity maps. More broadly, this strategy—training on physics-based dewetting free energies to build fast, context-aware predictors—is readily extensible to understanding and engineering systems where nanoscale surface solvation governs assembly, recognition, or material properties.

## Results

To establish ground truth for residue-level hydrophobicity, we first calculate the dewetting free energy, F_dewet_, around amino acids in different contexts using INDUS. In each case, we define a small observation volume enclosing the heavy atoms of the amino acid and compute the equilibrium work (i.e., free energy) required to remove water from that volume (Fig 1A). This construction lets us compare two contexts central to our study: isolated, capped amino acids in bulk water (Fig 1A) and amino acids embedded on protein surfaces with realistic curvature and chemical neighborhoods (Fig 1B). Across both settings, the INDUS protocol yields a direct, thermodynamic measure of how readily interfacial water dewets, providing a physically interpretable scale of context-aware hydrophobicity that we later use for model training and validation.

**Figure 1.**
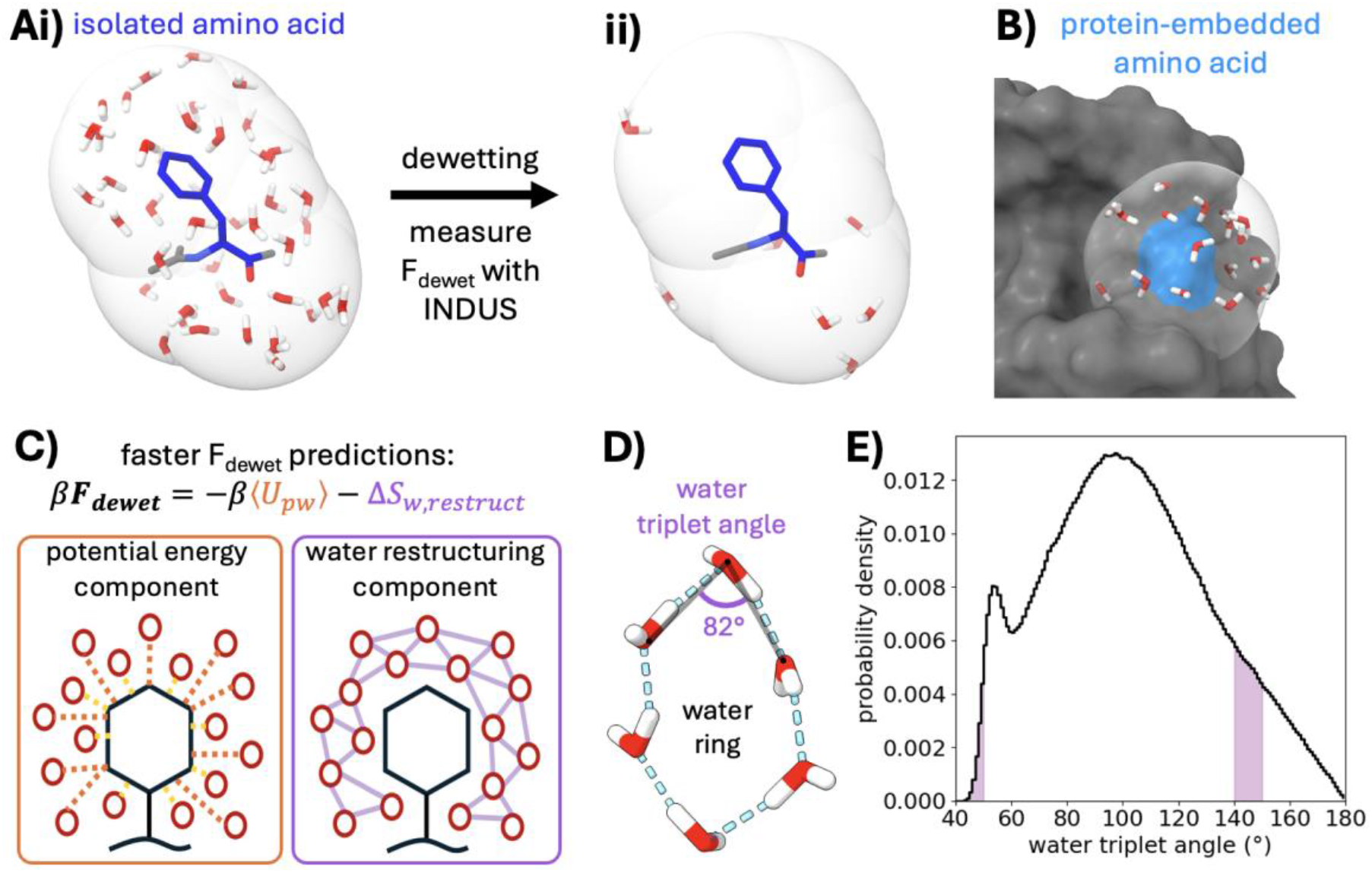
A) Dewetting free energies of isolated, capped amino acids are determined using the computationally intensive INDUS method, where umbrella sampling is used to evaluate the free energy difference between a fully solvated amino acid (i) and a dewetted amino acid (ii). Specifically, we measure the free energy to remove 80% of the solvation waters (i.e. waters within 5.5 Å of the amino acid’s heavy atoms, as indicated by the transparent shell). Previously we measured F_dewet_ for each isolated amino acid in three force fields. B) We additionally measured F_dewet_ of 40 amino acids embedded in structured proteins from hydrophobin and SARS-CoV-2 spike protein variants. Residue T28 from hydrophobin (blue) and its solvation waters are shown. C) A theoretical model predicts F_dewet_ from a short MD simulation by measuring the ensemble-averaged water-protein potential energies ⟨U_pw_⟩ and approximating the water restructuring entropy ΔS_w,restruct_ from water structural features. Water structural features include water triplet angle bins (D,E), water rings features (D), and H-bonding features.

Guided by a minimal physical picture (24, 43), we decompose *F*_*dewet*_ for an amino acid in water into a potential energy contribution from breaking residue–water interactions and an entropic contribution from releasing hydration-shell water. Hydration-shell water can have more pronounced structural correlations than bulk water, so its release upon dewetting is entropically favorable; this entropic gain competes against the energetic cost of severing residue–water contacts. We write this competition as:

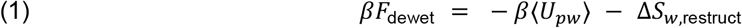

Here, ⟨*U*_*pw*_⟩ is the ensemble-averaged residue–water interaction potential energy; it is negative for favorable interactions, so − β⟨*U*_*pw*_⟩ > 0 penalizes dewetting. Δ*S*_*w*,restruct_ is positive by construction and captures the dimensionless entropy gained when hydration-shell water, initially ordered by the residue, is released and relaxes toward bulk structuring (such that −Δ*S*_*w*,restruct_ favors dewetting).β = 1/(*k*_*B*_*T*) is the inverse thermal energy. Hydrophobicity emerges when the entropic driving force exceeds the potential energy penalty.

This F_dewet_ formulation separates a directly measurable interaction term from a more elusive solvent entropy, motivating our approach for estimating each from simulation. ⟨*U*_*pw*_⟩ is obtained directly from a short, unbiased molecular dynamics simulation using interaction energy groups. We approximate the solvent restructuring entropy Δ*S*_*w*,restruct_ using water structural fingerprints and select them using a statistical learning approach, inspired by Dallin, et al. (30). Specifically, we assemble a feature pool of water fingerprints that includes water triplet angle distribution features (Fig 1E), water ring motifs (Fig 1D), and hydrogen-bonding features; all water fingerprints are measured within approximately two water layers of an amino acid. Our ultimate F_dewet_ models each include two water triplet angle distribution features, selected with LASSO, to complement ⟨*U*_*pw*_⟩ (see Appendix S1).

### HydroMap

We assemble two INDUS F_dewet_ datasets (Fig 2A). The isolated residues model set consists of 20 isolated amino acids with capped termini simulated in three widely used force fields (a99SB-disp, a03ws, and CHARMM36m) for 60 total conditions (42). The structurally-diverse model set consists of 40 protein-embedded amino acids from structured proteins (Table S1) alongside the 20 isolated amino acids (60 residues total in a99SB-disp), spanning diverse chemistries and local geometries.

**Figure 2.**
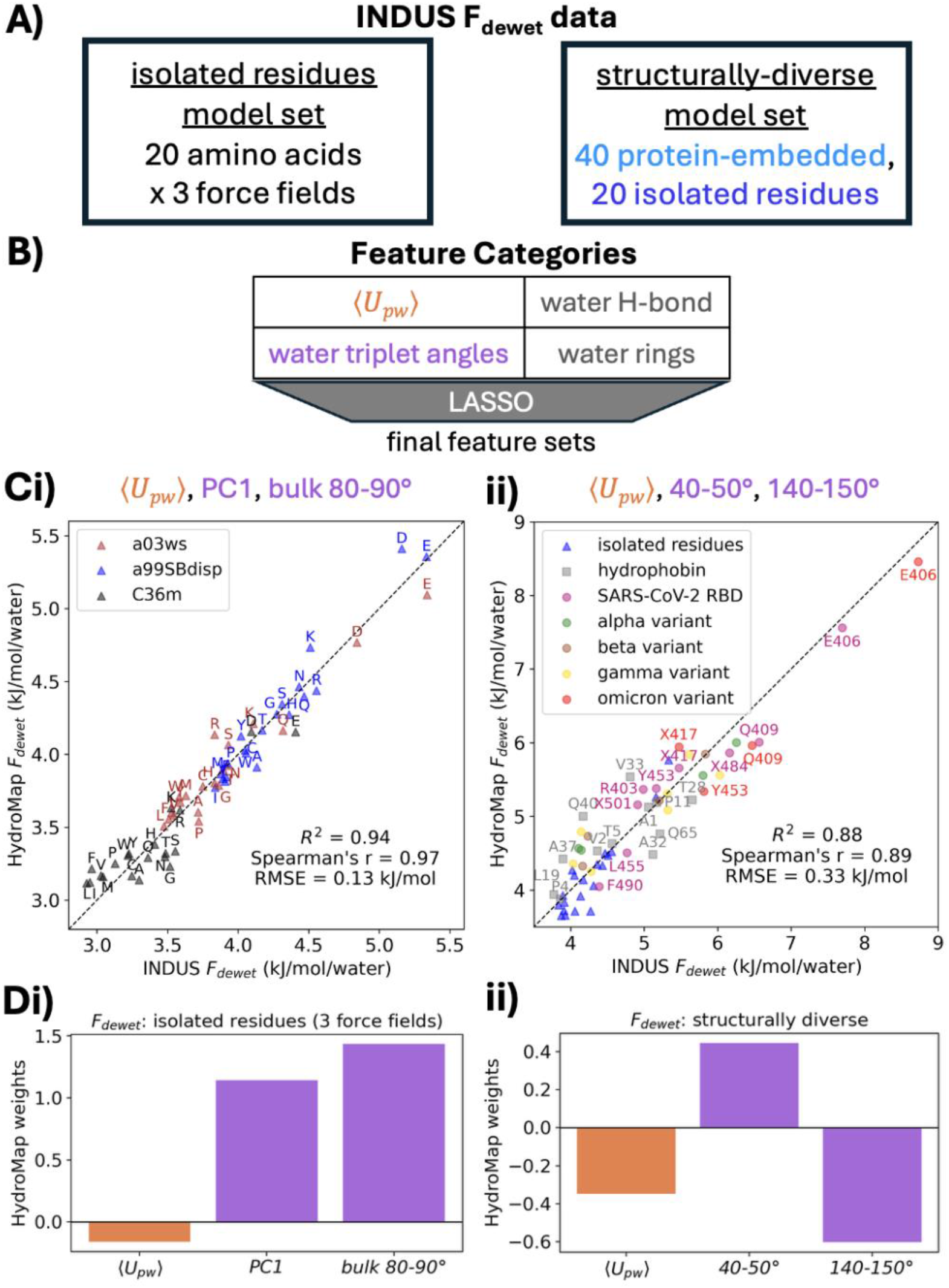
A) The INDUS F_dewet_ datasets consist of (i) the 20 capped amino acids simulated in three force fields (a99SB-disp, a03ws, and CHARMM36m), and (ii) 40 amino acids embedded in a protein context from structured proteins (from hydrophobin and SARS-CoV-2 variants) simulated in a99SB-disp. B) The feature pool for the linear HydroMap models consists of ⟨U_pw_⟩, water triplet angle features (including 10° bins and principal component loadings), water hydrogen bond features, and water ring features (see Appendix S1); up to two features were selected with LASSO to complement ⟨U_pw_⟩ in these models. C) Two features—⟨U_pw_⟩ and water triplet distribution’s PC1—achieved R^2^ = 0.94 for one force field (a99SB-disp), and adding bulk water triplets’ 80-90° frequency (approximating bulk water entropy) enabled the models to achieve R^2^ = 0.94 across isolated amino acids in all three force fields. D) Three features—⟨U_pw_⟩, and the water triplet distribution’s 40-50° and 140-150° angle frequencies—achieved R^2^ = 0.88 for the 40 protein-embedded amino acid dataset, with nearly identical performance when including the also the 20 isolated amino acids. The weights for the standardized features of the two main models are shown in (Cii) and (Dii).

### Isolated residues simulated in a99SB-disp

We observe that a two-feature model, consisting of ⟨*U*_*pw*_⟩ and a water triplet angle feature called PC1, closely recovers INDUS F_dewet_ on the twenty isolated, capped residues simulated in the a99SB-disp force field (R^2^ = 0.94; RMSE = 0.09 kJ/mol per water). The PC1 feature is a measure of the primary mode by which water’s 3-body angle correlations change (25), and is physically interpretable as a tradeoff between simple fluid-like (i.e. more frequently icosahedral) and water-like (i.e. more frequently tetrahedral) hydration water angle structure; see Fig S1. Generally, isolated amino acids with more tetrahedral water structuring (i.e. negative PC1 values) are more hydrophobic (i.e. lower F_dewet_).

### Small hydrophobe dataset: isolated residues simulated in three force fields

Extending this model to the three-force field set requires one additional descriptor to accommodate differences in bulk water structuring: adding the bulk 80–90° triplet-angle fraction yields R^2^ = 0.94 (RMSE = 0.13 kJ·mol^−1^ per water) across all 60 isolated amino acids (Fig 2B i). We interpret this bulk 80–90° triplet-angle as correcting for differences in neat water configurational entropy between force fields. In an ideal gas—with no structuring and high configurational entropy—the triplet angle distribution follows *sin(θ)*. CHARMM36m more closely resembles the *sin(θ)* distribution while a99SB-disp and a03ws have broad increases in structuring around 80-125° (Fig S2-3). The bulk 80-90° triplet angle is therefore correlated with greater deviations from ideal gas behavior and therefore reduced configurational entropy. Dewetting is hindered by reduced bulk water entropy, since there is a reduced entropic driving force (ΔS) between the residue’s ordered hydration waters (low S) and the disordered bulk (higher S). Accordingly, a larger bulk 80–90° fraction—signaling more structured, lower-entropy bulk water—shrinks the ΔS between bulk water and the ordered hydration shell, thus opposes dewetting and yielding a positive association with F_dewet_ across force fields (Fig 2B ii).

### Structurally diverse dataset: Protein-embedded and isolated residues

Next, we extend our analysis to protein-embedded residues and find that a sparse model with just three terms—⟨*U*_*pw*_⟩ and the 40–50° and 140–150° triplet-angle bins—captures INDUS *F*_dewet_with high accuracy (*R*^2^= 0.88; Spearman *r* =0.89; RMSE =0.37kJ·mol^−1^ per water). This performance is essentially unchanged when the model is fit jointly with the isolated-residue data (*R*^2^= 0.88; *r*=0.89; RMSE =0.33kJ·mol^−1^ per water), as seen in Figure 2C i.

The hydrophilic 40-50° feature increases with highly coordinated waters at polar and charged sites, consistent with prior observations of over-coordinated waters at hydrophilic sites (26, 30, 44). This feature is associated with higher F_dewet_ values (hydrophilicity) and corresponds with PC3, the third principal component of the water triplet distribution (Fig S1). By contrast, the 140-150° feature is a hydrophobic signature. The 140-150° water triplet population is generally highest in bulk water and in the isolated amino acid dataset (i.e. small hydrophobes). In structured proteins, a number of convex residues also had relatively large 140-150° water triplet populations, such as A1 & V2 in hydrophobin II and a solvent-exposed F490 in SARS-CoV-2’s receptor binding domain (Fig S4). Meanwhile some protein-embedded residues in flat/concave geometric contexts have strong depletions in 140-150° angles, such as P11 in hydrophobin II and E406 in SARS-CoV-2’s receptor binding domain (Fig S4). We observe a similar pattern in a α-synuclein amyloid filament where a surface-exposed alanine in the middle of a filament (flat/concave environment) has a depleted 140-150° water triplet population relative to the same alanine in more convex environments (Fig S5).

To further dissect the topological drivers of this 140-150° angle depletion, we examined how curvature alone modulates the 140–150° population. We find that surface curvature has a strong, monotonic influence (Fig S6): increasingly flat interfaces show progressively greater depletion of the hydrophobic 140–150° triplet signature. Even hydrophobic surfaces like hard spheres/cubes can induce this hydrophilic depletion of the 140-150° triplet signature (Fig S6). Furthermore, across the entire dataset, the 140-150° signature appears to be strongly anti-correlated with PC2 (r = -0.84), which balances ideal-gas-like 90° angle populations against large-angle and icosahedral motifs (Fig S1) and features prominently around extended hydrophobic surfaces, such as methylated SAMs (25). Interestingly, we see strong correlations to certain hydrogen-bonded water ring features (r = 0.74-0.78 for 4-6 membered rings; Fig S7); no other 10° water triplet bin is more correlated with the presence of 4-6 membered water rings. These observations suggest that local topology is a key determinant in 140-150° depletion—a phenomena associated with hydrophilicity.

Together, these HydroMap results show that residue–water potential energy and two water structural fingerprints largely explain *F*_dewet_ across isolated and embedded contexts with a variety of chemical and geometric environments. In addition, a single bulk water structural feature corrects for hydrophobicity shifts between force fields by approximating bulk water entropy differences. With these simple, interpretable HydroMap models in hand, we can rapidly predict F_dewet_ at the scale of thousands of residues by running short MD simulations and measuring residue–water potential energy and water structure (Fig 3A).

**Figure 3.**
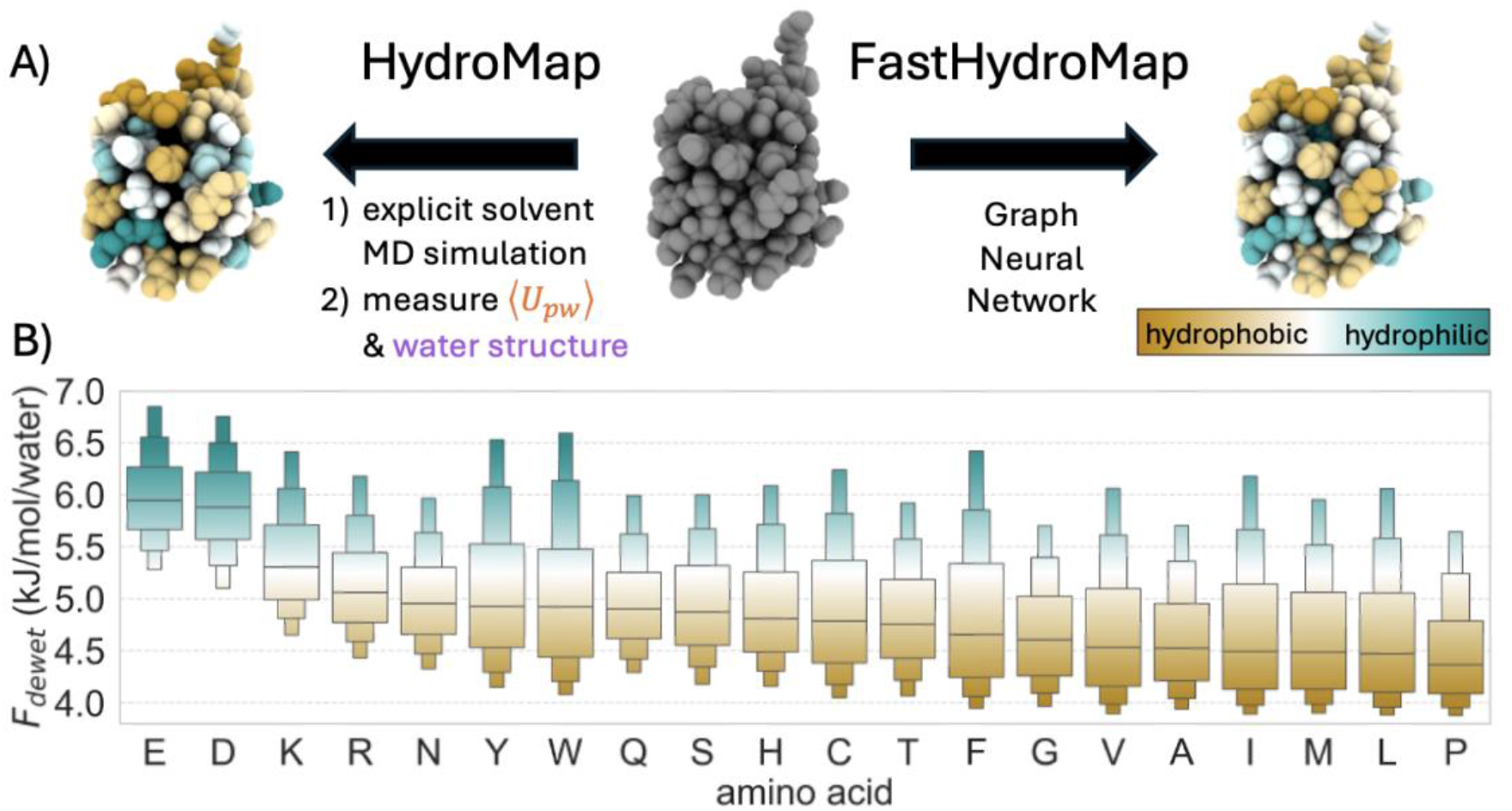
HydroMap and FastHydroMap for residue-level hydrophobicity estimation. A) Schematic of the HydroMap workflow, which uses explicit-solvent molecular dynamics simulations to compute residue–water interaction energies ⟨U_pw_⟩ and local water structure, enabling prediction of residue-level dewetting free energy (F_dewet_), for ubiquitin in this case (PDB #1UBQ). FastHydroMap replaces this explicit solvent simulation pipeline with a graph neural network (GNN) that predicts F_dewet_ directly from protein structure, enabling rapid mapping of hydrophobic (gold) and hydrophilic (teal) regions. B) Distributions of F_dewet_ values of 20 amino acid types from applying HydroMap to 931 protein structures from the PDB, totaling ∼66.8 k solvated residues. Boxenplots show the F_dewet_ distributions with three levels, e.g. showing the 50^th^, 75^th^, 87.5^th^, and 93.75^th^ percentiles. The twenty amino acids are ordered by their median F_dewet_ from hydrophilic to hydrophobic.

### FastHydroMap

While HydroMap is much faster than explicit INDUS calculations, it requires short explicit-solvent MD simulations to evaluate the residue–water potential energy and water structural metrics. To remove the need for simulations, we develop FastHydroMap, a lightweight graph neural network trained to reproduce HydroMap residue labels directly from protein structure. FastHydroMap represents a protein as a residue graph and predicts residue-level F_dewet_ from amino-acid identity, solvent exposure, and local geometry. In FastHydroMap’s architecture (Fig 4), each residue’s F_dewet_prediction is decomposed into an intrinsic term computed from residue features before message passing and a contextual term computed after message passing, allowing the model to separate local chemistry from neighbor-dependent effects.

**Figure 4.**
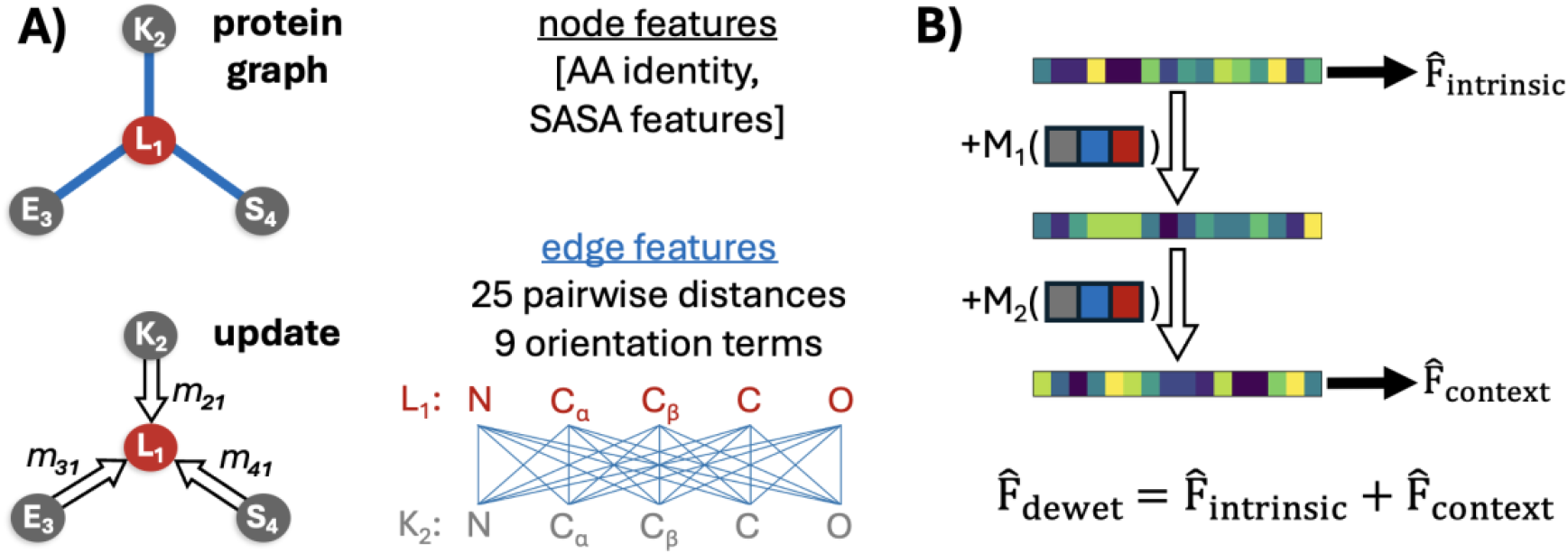
FastHydroMap architecture. A) The protein backbone is represented as a graph where each residue is a node connected to its nearest spatial neighbors. Node features encode amino acid identity and solvent-accessible surface area; edge features encode 25 pairwise distances between backbone heavy atoms (N, Cα, C, O, and Cβ), expanded in a radial basis, plus 9 terms capturing the relative orientation of neighboring backbone frames. During message passing, each residue updates its representation by aggregating learned messages from its neighbors, incorporating local structural context. B) The model predicts the dewetting free energy (F_dewet_) as a sum of an intrinsic term, 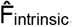, derived from per-residue features, and a context-dependent term, 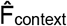, computed after message passing. This decomposition expresses the physical intuition that each amino acid has a baseline dewetting propensity that is modulated by its surrounding environment. See Methods and Appendix S2 for full details.

We generate FastHydroMap training data by applying HydroMap to 931 curated proteins, yielding 66,827 well-solvated residues for evaluation. The resulting F_dewet_ distributions span broad ranges within each amino acid type (Fig 3B), especially for aromatic residues. For example, the bottom quartile of solvated tyrosine residues in our dataset have F_dewet_ < 4.5 kJ·mol^−1^ per water while the top 12.5 percent have F_dewet_ > 6 kJ·mol^−1^ per water. Residues of identical chemical identity can occupy markedly different solvation environments, underscoring a central limitation of hydropathy scales that assign a single value to each residue.

FastHydroMap is a compact message-passing network with only 46,474 parameters. Model architecture and hyperparameters are selected on the validation set, and the held-out test set is used only for final evaluation. On that held-out set, the model achieves an RMSE of 0.472 ± 0.01 kJ mol^−1^ per water (Pearson r = 0.788 ± 0.001, Spearman ρ = 0.790 ± 0.0003, averaged over three random initializations). Ablations on the validation set reveal that local residue features — amino acid identity and solvent exposure — capture much of the HydroMap signal (see Appendix S2). Message passing provides a consistent additional improvement. The mean absolute correction of the contextual F_dewet_ term is ∼0.22 kJ mol^-1^ per water, with approximately 10% of residues having a contextual F_dewet_ term of greater than 0.44 kJ mol^-1^ per water. The same architecture also predicted water triplet-angle principal components (PC1, PC2, PC3) from the protein structure with a held-out Spearman ρ = 0.77 –0.80 (Fig S8).

FastHydroMap is fast enough for routine large-scale use on ordinary CPUs. On a desktop Intel i9-14900K CPU, end-to-end inference took ∼0.08 s for a 56-residue protein, ∼1.3 s for a 730-residue multichain assembly, and ∼7.6 minutes for a 13,000-frame trajectory of a 56-residue protein (∼35 ms per frame). These speedups make it practical to score every residue across protein trajectories and to support high-throughput tasks such as mutational scanning or surface redesign, without explicit-solvent simulation. We next illustrate these capabilities in a series of case studies.

### Case Studies

#### Parkinson’s amyloid filament hydrophobicity

We first apply HydroMap to the cryo-EM structure of the Parkinson’s disease filament (PDB #8A9L). This structure consists of stacked α-synuclein protofilaments along with two unassigned linear peptide densities that thread along two distinct interfaces of the α-synuclein core, near residues 50-55 and 84-91 (Fig 5A i). Classical hydrophobicity scales, such as the MLP hydrophobicity scale, characterize these interfaces as only weakly hydrophobic and do not identify them as favorable dewetting sites (Fig 5B). In contrast, HydroMap’s context-aware F_dewet_ map reveals both interfaces as highly hydrophobic and thus favorable dewetting locales. The high hydrophobicity (low F_dewet_) at these two interfaces indicates a strong thermodynamic driving force for water exclusion upon binding and is consistent with the preferential adhesion of the unassigned peptide densities to these positions. Notably, both interfaces exhibit pronounced tetrahedral water structuring (see PC1 map in Fig S9), consistent with the presence of highly ordered, low-entropy interfacial waters. Displacement of these waters upon peptide binding provides a substantial entropic gain, directly lowering F_dewet_ and generating an effective hydrophobic driving force that is not captured by classical static hydrophobicity scales.

**Figure 5.**
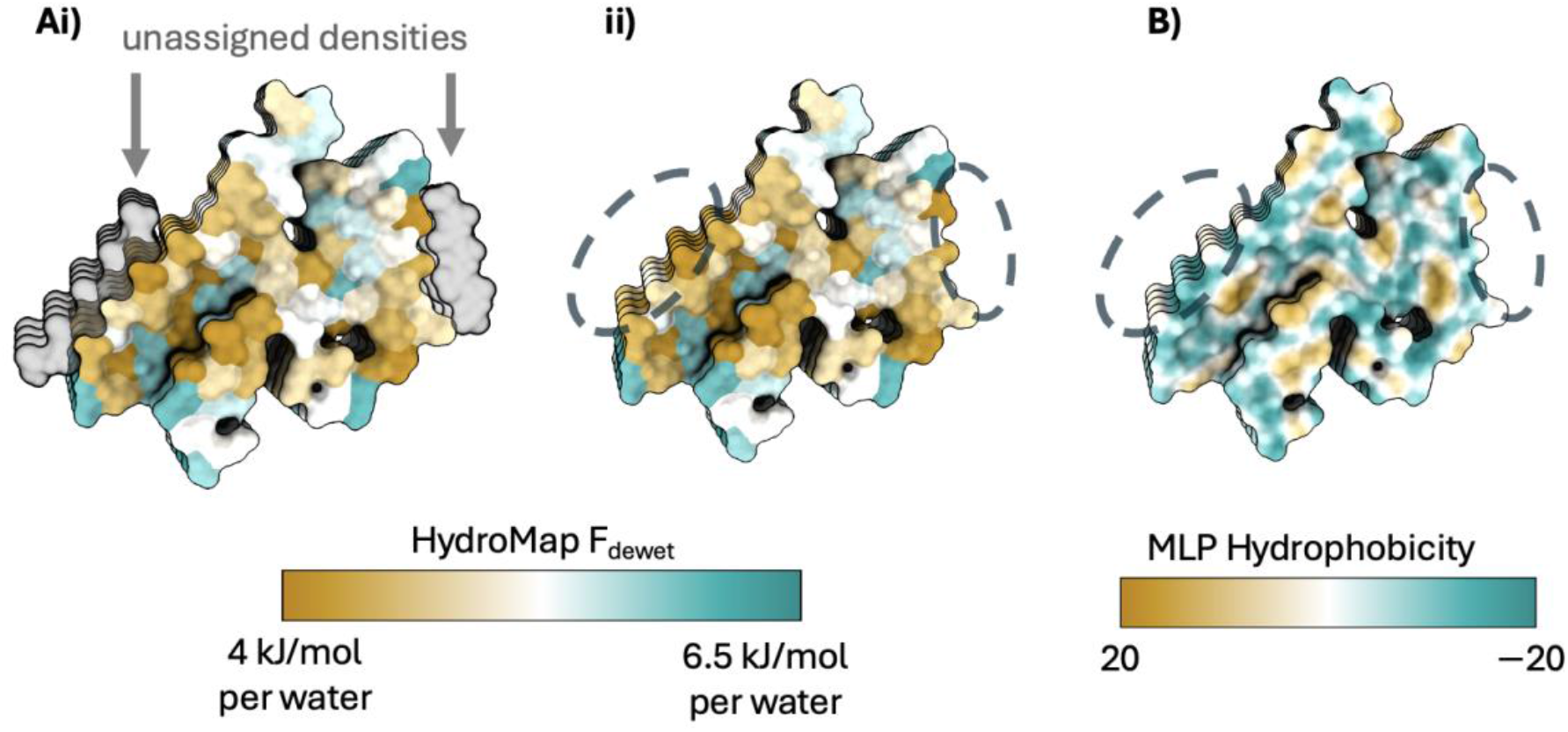
A) The cryo-EM structure of the α-synuclein amyloid filament from Parkinson’s disease patients (PDB #8A9L) contains two unassigned linear peptide densities: one near residues 84–91 (left, gray density) and another near residues 50-55 (right, gray density) as seen in panel (i). (ii) Coloring by HydroMap-predicted dewetting free energy (F_dewet_) reveals that both interfaces are strongly hydrophobic. B) In contrast, these hydrophobic interfaces are not signaled by the molecular lipophilicity potential (MLP) scale, highlighting the inability of traditional hydrophobicity metrics to capture interfacial water thermodynamics driven by local geometry and chemistry.

#### Calmodulin: a fold-switching calcium sensor

We apply FastHydroMap to calmodulin, a Ca^2+^ sensor that toggles between an apo and Ca^2+^-bound state. Calmodulin has EF-hand motifs—helix-loop-helix structures that each coordinate a Ca^2+^ ion. Ca^2+^ binding reorganizes a methionine-rich region into a helix-binding groove. Applying FastHydroMap to both crystal structures and focusing on the C-terminal lobe, we observe a concerted increase in hydrophobicity across groove residues I85, A88, M109, L112, I130, M144, M145, and T146 (Fig 6A; Fig S10). These hydrophobicity shifts are not visible to static scales. For most groove residues this hydrophobicity shift reflects changes in solvent exposure (captured by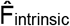, but for residues like I85 and M109 it is driven primarily by changes in neighboring residues (captured by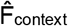; see Fig S11.

**Figure 6.**
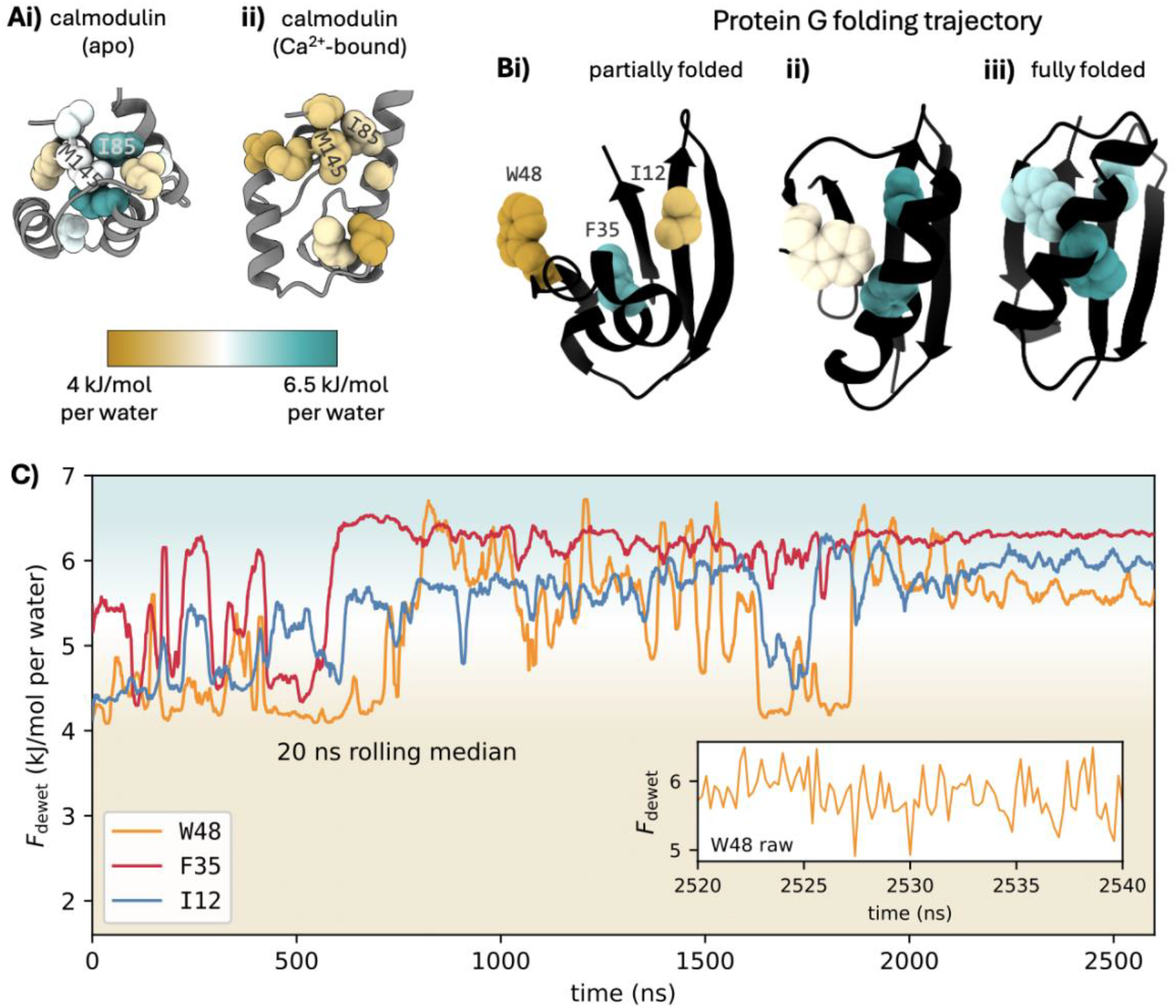
A) Calmodulin undergoes a Ca^2+^-induced conformational transition from a closed (PDB #1CFD) to an open state (PDB #1CLL), accompanied by a decrease in local dewetting free energy (F_dewet_) at a helix-binding interface. B) Representative snapshots of a folding trajectory of Protein G’s GB1 domain illustrate how residue-level hydrophobicity evolves during folding. The residues W48, F35, and I12 exhibit distinct changes in Fdewet as the protein transitions from partially folded to fully folded conformations. C) Time series of Fdewet for W48, F35, and I12 over the full folding trajectory reveal fluctuations on multiple timescales, including tens-to-hundreds of nanoseconds associated with major conformational rearrangements and nanosecond-scale fluctuations within a single folded basin (inset). A 20 ns rolling median is plotted in panel C for visual clarity, while the inset plots raw F_dewet_values. Residue-level F_dewet_ values for the 56 GB1 residues in each frame of a 13,000-frame MD simulation were computed by FastHydroMap in ∼7.6 minutes on an Intel Core i9-14900K CPU and ∼14.1 minutes on a Macbook Pro with an Apple M1 chip.

#### Protein G: dynamic hydrophobicity in a protein folding trajectory

Finally, we apply FastHydroMap to a long molecular dynamics folding trajectory of the GB1 domain of Protein G, a well-characterized system for protein folding (45–51), to visualize the hydrophobic effect in action at residue resolution. We analyze a 2.6 μs (13,000-frame) trajectory from DE Shaw Research (49) of the 56-residue NuG2 variant (50), predicting F_dewet_ for each residue in each frame. This analysis makes it possible to follow how local hydration environments reorganize as folding proceeds. We find that, in partially folded conformations, some solvent-exposed patches are strongly hydrophobic (low F_dewet_). Then as the protein folds, these patches are buried or reorganized, and the solvent-facing surface becomes enriched in more hydrophilic surfaces (high F_dewet_).

The time series further show that this hydrophobic surface reorganization occurs across multiple timescales. Residues such as W48, F35, and I12 undergo large increases in Fdewet over hundreds of nanoseconds as the trajectory commits toward the folded basin (Fig 6C, Movie S1). When we compare the unfolded and folded conformational ensembles, we find that the largest upward F_dewet_ shifts are concentrated in the conserved GB1 folding nucleus identified by prior simulation and sequence-conservation analyses (Fig S12) (52, 53). That a hydration-based metric independently highlights these residues is suggestive that F_dewet_ reports on the same underlying property that makes them evolutionarily preserved and kinetically important. A minority of residues shift in the opposite direction; residue T54 shows the largest such shift (Fig S12).

Superimposed on these slow, concerted motions, residues show nanosecond-scale fluctuations in F_dewet_. We focus on the F_dewet_ fluctuations of W48 and how it appears to have a particularly varied solvation environment, indicated by its F_dewet_ distribution, even in the folded confirmation (Fig 6C inset, Fig S13). Residue-level hydrophobicity is not a fixed property of the native state, but a dynamic field that continues to fluctuate around the folded structure. These context-dependent, dynamic shifts in hydrophobic driving forces are invisible to the classic and widely referenced hydrophobic–polar (HP) model of folding, which assigns each residue a fixed hydrophobicity label. Here, the revealed dynamic hydrophobic forces have remarkably interesting potential implications for how hydrophobic forces dynamically shape folding pathways and mechanisms, and how protein structures and sequences may have evolved to utilize them. FastHydroMap’s speed—predicting F_dewet_ for all 56 residues across 13,000 frames in ∼7.6 min (see Fig 6 caption)—makes this kind of time-resolved revelatory analysis possible for folding trajectories, conformational transitions more broadly, and other high-throughput applications.

## Discussion

Our results suggest that residue-level dewetting free energies in proteins can be inferred with high accuracy from statistics of the fully solvated state. While dewetting is a property of rare, low-density water fluctuations, it appears that dewetting propensity can be predicted with less water sampling than expected: just a few hundred picoseconds of sampling per solute conformation (Fig S14). By combining residue-water interaction potential energy ⟨*U*_*pw*_⟩ with two local water-structure fingerprints, HydroMap recovers INDUS F_dewet_ for both isolated and protein-embedded amino acids across diverse chemical and geometric environments. The functional form of HydroMap is motivated by a simple solvation-thermodynamics picture (Eq 1), in which removing water incurs a potential energy penalty from breaking residue–water interactions, while local water-structural signatures serve as proxies for the entropic cost or benefit of solvent restructuring. In this respect, the success of HydroMap’s prediction abilities points to the utility of this simple decomposition of solvation free energy into direct interactions and water restructuring entropies.

Specific features of the triplet-angle distribution (PC1, and 40–50° & 140–150° bins) emerge as robust correlates of F_dewet_, meaning that the local structural organization of water hydration network around a residue encodes its propensity to dewet even when no dewetting event has occurred. Notably, these 40-50° and 140-150° triplet angle descriptors are consistently selected over alternative water-structure fingerprints (including hydrogen-bond and water-ring metrics). PC1 captures the balance between tetrahedral and icosahedral water motifs, and we find that more tetrahedral local water structure generally accompanies more favorable (lower) F_dewet_ in small hydrophobes (e.g., isolated amino acids). The 40–50° bin is enriched at hyper-coordinated waters near charged and strongly polar sites, and correlates with hydrophilicity and higher F_dewet_. Conversely, the 140–150° bin is a signature of small hydrophobes and bulk-like water, and its depletion appears to coincide with surface curvature (Fig S4-6) and other topological and chemical environments that resist dewetting (i.e. hydrophilic surfaces). Together with ⟨U_pw_⟩, these features can be viewed as an approximate decomposition into a term dominated by residue–water interaction energies and a term that largely reflects solvent restructuring, ΔS_w,restruct_.

Our analysis across three force fields further highlights the thermodynamic nature of these descriptors. A single bulk-water triplet feature (the 80–90° fraction) compensates for systematic shifts in F_dewet_ between force fields (Fig S2-3, Fig S15), consistent with the idea that differences in bulk water entropy modulate the driving force for expelling ordered hydration waters. In other words, much of the force-field dependence of F_dewet_ may be due to the contrast in entropy between bulk water and the hydration shell. This perspective provides a simple lens for comparing hydrophobicity across force fields: water models with more structured, lower-entropy bulk water (reflected by 80– 90° fractions) naturally show larger dewetting costs, while models with more disordered bulk water exhibit a stronger hydrophobic driving force (e.g., CHARMM36m (42, 54)). In principle, such bulk-structure observables could even be used as a tuning knob when assessing or refining force fields for interfacial hydrophobicity.

### Hydrophobicity as contextual and dynamic

By training HydroMap and FastHydroMap on diverse chemical and geometric environments, we directly observe the strongly contextual nature of hydrophobicity. Each amino acid type spans a broad distribution of F_dewet_ values across protein structures (Fig 3A), particularly aromatics such as F, Y, and W, which can range from highly hydrophobic to relatively hydrophilic depending on their local topology, neighbors, and accessibility. Static hydrophobicity scales that assign a single value per residue cannot represent this diversity. In contrast, residue-resolved F_dewet_ accounts for nuances in water restructuring and thus is sensitive to conformational state and other subtle differences in the microenvironment.

FastHydroMap’s speed makes it practical to evaluate hydrophobicity across the full conformational ensemble that a molecule explores. Such time- and ensemble-resolved analyses are largely inaccessible to rigorous INDUS F_dewet_ calculations, which require extensive sampling of water-density fluctuations for each conformation of interest. Proteins and other macromolecules undergo both local and global conformational changes over a range of timescales, and the hydrophobicity of individual residues can shift substantially as side chains rearrange, loops fluctuate, or domains move. Because FastHydroMap rapidly predicts F_dewet_ directly from protein geometry, it can track hydrophobicity along large-scale conformational transitions and folding trajectories (Fig 6). In calmodulin, we see a concerted increase in hydrophobicity across the helix-binding groove upon Ca^2+^ binding, driven purely by geometric rearrangement. In Protein G, we observe slow, cooperative shifts in hydrophobicity as residues become buried or reoriented during folding, along with rapid nanosecond-scale fluctuations driven by side-chain motions. These examples emphasize that hydrophobicity is not a static property of residues but a dynamic field on the protein surface that evolves with structure and time. The implication is that dynamic hydrophobic driving forces shape these conformational pathways, not merely the static hydrophobic-hydrophilic patterning as has long been used as a basic biophysical model for protein folding.

### INDUS to HydroMap to FastHydroMap: a general approach

Beyond the specific models presented here, our approach offers a generalizable strategy for turning expensive thermodynamic calculations that sample rare states into practical, interpretable, and eventually ultrafast predictors. First, we perform a limited number of rigorous INDUS calculations, to generate high-quality F_dewet_ labels for a carefully chosen set of environments. Second, we build a sparse, physics-informed model (HydroMap) that expresses F_dewet_ in terms of ⟨U_pw_⟩ plus a small number of water-structure fingerprints, ideally features that are physically interpretable. Third, we train an ultrafast model (FastHydroMap) that can use graph neural networks (or other ML approaches), to reproduce the F_dewet_ predictions with a lower computational cost, eliminating the need for solvent simulations altogether. This three-tiered approach provides rigorous reference data, reveals governing features, and operationalizes them for large-scale use; researchers can then pick a model that fits a desired speed–accuracy tradeoff. In practice, HydroMap achieves an RMSE of 0.33 kJ/mol per water against INDUS, and FastHydroMap trades a modest accuracy cost (0.47 kJ/mol per water) for orders-of-magnitude faster inference.

This three-tiered strategy is broadly applicable to other solvation-related observables that are currently accessible only through enhanced sampling, including cavity formation free energies in materials or wet-dry transitions in pores. HydroMap was trained on 20 canonical amino acids that span a wide chemical space; we expect it to provide a reasonable first approximation of desolvation free energies for many environments out-of-the box. When more accuracy is required or when new chemistries are introduced (e.g. noncanonical amino acids, polymers, peptoids, nucleic acids, etc.), additional INDUS calculations can be included into the training sets to refine both HydroMap and FastHydroMap. Additionally, one could also enrich the feature set by including descriptors measured in partially desolvated simulations—e.g., sampling both the unbiased state and a modestly dehydrated state. This would double the simulation cost per site, but could expose additional fingerprints of the wet–dry transition and further sharpen linear models.

### Biological implications and practical applications

The case studies above illustrate two complementary uses of context-aware F_dewet_: revealing hidden structural features on static surfaces, and tracking how hydrophobicity evolves with conformational changes. In the α-synuclein amyloid filament associated with Parkinson’s disease, HydroMap identifies two strongly dewetting interfaces that coincide with unassigned peptide densities running along the fibril surface. Classical hydropathy metrics fail to flag these sites as particularly special. HydroMap’s F_dewet_, by contrast, recognizes them as thermodynamically poised for water exclusion, consistent with their apparent role in recruiting the unknown peptide. More generally, context-aware F_dewet_ can reveal “hidden” binding sites on amyloid or protein surfaces that do not stand out in contextless hydrophobicity scales.

The calmodulin and Protein G case studies underscore the utility of time-resolved F_dewet_. In calmodulin, the Ca^2+^-induced opening of the EF-hand motifs yields a coordinated increase in hydrophobicity at the helix-binding groove, aligning with its known role as a promiscuous binding platform. In Protein G, by scoring F_dewet_ along a long folding trajectory, we directly observe the hydrophobic effect in action: solvent-exposed dewetting-prone patches are systematically replaced by more water-stabilizing exposure as folding proceeds. From this perspective, folding is not simply a matter of burying hydrophobic residues, but a coordinated restructuring of the protein surface that progressively stabilizes water. Many residues whose hydrophobicity shifts most strongly during folding are the same ones independently identified as folding-nucleus members or as evolutionarily conserved sites (Fig S12) (52, 53), suggesting that F_dewet_ reports on the underlying property that makes these residues mechanistically central. Taken together, these examples highlight how a single, residue-resolved wetting metric can connect angstrom-scale hydration structure to emergent interface properties that matter at the nanometer scale in binding, assembly, and folding. More broadly, these F_dewet_ models may enable more sophisticated folding and conformational sampling models that treat the solvent as an active participant rather than an implicit background. Proteins fold differently—or fail to fold—in modified solvents, emphasizing that water’s local structure and fluctuations shape the free-energy landscape in ways that can depend subtly on chemical and geometric context. Many implicit-solvent treatments collapse these effects into coarse surface-area related terms and therefore may miss nuances in hydration that distinguish dewetting-prone from water-stabilizing surfaces. In contrast, F_dewet_ is defined as a free-energy cost of removing waters from a specified volume, and HydroMap/FastHydroMap make that thermodynamic quantity practical to compute at the residue level.

In principle, this makes F_dewet_ a natural candidate for inclusion in energy and scoring functions across a range of structural modeling tasks. In Monte Carlo protein folding or conformational sampling schemes, proposed moves (e.g., side-chain rotations, backbone pivots, domain rearrangements) can be accepted or rejected based in part on changes in F_dewet_. F_dewet_ could likewise enter the scoring and refinement stages of AI-based structure prediction pipelines, where physically grounded solvation terms may help discriminate among candidate conformations that score similarly under learned potentials. While protein folding remains challenging, particularly in regimes where evolutionary information is sparse, improving energy functions with efficient, context-aware hydration physics may help guide sampling toward more realistic conformations.

More fundamentally, physics-based modeling frameworks share a common limitation in how hydrophobicity itself is represented. HP models and many coarse-grained potentials assign each residue a fixed hydrophobicity label, independent of orientation or environment. Implicit-solvent treatments are more sophisticated but share the underlying assumption: hydrophobic effects are typically collapsed into surface-area-based terms or generalized solvation energies that do not capture the geometry-dependent water restructuring that drives them. This approach works on average but cannot reproduce the strong context-dependence of hydrophobicity documented above. Rapid F_dewet_ predictions offer a path forward: fixed hydrophobicity labels and surface-area-based solvation terms can be replaced with context-aware driving forces grounded in explicit-solvent physics. Looking further ahead, future force field architectures may separate out geometry-dependent hydration physics from dispersion forces, rather than absorbing them into a single tuned interaction.

Beyond improving physics-based modeling, we envision F_dewet_ maps as a routine tool for protein design and mechanistic interpretation. For example, protein design may use F_dewet_ predictions to engineer dewetting-prone patches to strengthen binding interfaces, or to engineer dewetting-resistant surfaces to reduce nonspecific adhesion and unwanted aggregation. In conformational dynamics and signaling, large shifts in F_dewet_ can help identify key surfaces for binding or folding that would be missed by static, contextless hydrophobicity scales. Finally, binding thermodynamics often hinge on desolvation (two partners must dewet before binding), and F_dewet_ provides a physically grounded complement to commonly used implicit solvation energies. In practice, FastHydroMap makes it feasible to evaluate F_dewet_ across large panels of candidate binders or interface designs, and to incorporate solvation energy terms alongside other more accessible energy contributions. In this way, F_dewet_ offers a practical bridge between detailed explicit-solvent physics and the heuristic energy models that currently drive most protein engineering workflows.

### Outlook

HydroMap and FastHydroMap predict F_dewet_ in a substantial desolvation regime that characterizes binding and folding processes, where surfaces often do not dry out completely. The same INDUS with machine learning approach generalizes naturally across the full range of water exclusion, from the shallow desolvation in weak or transient interactions to near-complete desolvation in tightly packed interfaces; the water-structure features most predictive of F_dewet_ may shift across this range. Generating training data at additional extents of water removal, or predicting the entire dewetting free energy profile (10) rather than a single value, is a natural extension and would map how hydration physics varies with the depth of water exclusion.

The training set used here emphasizes small, globular proteins in aqueous solution, which span the regimes most relevant to the case studies presented. Extending the framework to less-solvated environments, such as deeply buried cavities and pores, is a promising direction and will benefit from targeted INDUS calculations in these regimes. The extent to which residue-level F_dewet_ transfers to larger patches, or whether collective hydration effects at the patch scale produce non-additive behavior, is an open question of direct relevance to interface dewetting. Extending the framework to patch-resolved F_dewet_ is a natural way to test this. Beyond proteins, the same approach extends naturally to other biomolecular and material surfaces — including nucleic acids, lipid bilayers, glycans, and synthetic polymers — wherever local solvation governs assembly, recognition, or transport. Dewetting depth, environmental scope, spatial resolution, and chemical class are each interesting directions in which the framework can extend, each tied to distinct questions in hydration physics.

## Conclusion

Hydrophobicity is a contextual, dynamic property rather than a fixed label of amino acid identity. Dewetting is governed by rare fluctuations in water density, yet our results show that its thermodynamic signature can be inferred from statistics of the fully solvated state — revealing that local hydrophobicity is encoded in a small number of low-order structural features of interfacial water, and that residue-level dewetting free energies can be predicted without explicitly sampling low-density states. HydroMap and FastHydroMap translate this physical picture into practical tools, enabling routine evaluation of Fdewet in static structures and trajectories. Applied to biological systems, they expose dewetting-prone interfaces on amyloid filaments that contextless hydrophobicity scales miss, resolve a concerted hydrophobic shift across calmodulin’s helix-binding groove upon Ca^2+^ binding, and track the hydrophobic effect residue-by-residue through a Protein G folding trajectory.

More generally, this work illustrates how rare-event thermodynamic observables can be approximated, interpreted, and accelerated to bridge molecular physics and high-throughput modeling. By making HydroMap and FastHydroMap openly available, we aim to facilitate broader use of dewetting-based hydrophobicity in biological and materials systems, and to support folding, binding, and design workflows that treat solvent as an active, context-sensitive participant in the energy landscape.

## Methods

### INDUS training-label generation

Residue-level F_dewet_ labels for HydroMap were generated using indirect umbrella sampling (INDUS), following our prior amino-acid dewetting protocol. INDUS simulations were performed with a modified version of GROMACS 2016.3 in which a harmonic bias acts on the coarse-grained number of water molecules *N*_*v*_ inside a residue-centered hydration volume. The biased system is governed by the Hamiltonian 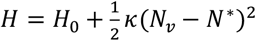, where *H*_0_ is the unbiased potential, *N*^*^is the umbrella center, and the bias strength is κ = 10 κ_0_ with 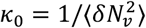 the inverse variance of unbiased water-number fluctuations. For each residue, this volume was defined as the union of spheres of radius *R*_*v*_ =0.55nm centered on the residue’s heavy atoms, corresponding approximately to waters within 5.5 Å of the residue. During these INDUS calculations, the residue heavy atoms were restrained to preserve the target local geometry while waters were biased out of the observation volume. The Gaussian coarse-graining parameters were σ=0.01nm and *r*_*c*_ = 0.02 nm. Overlapping *N*_*v*_ windows were simulated for 6 ns each, discarding the first 2 ns as equilibration, and the unbiased *P*_*v*_(*N*)distribution was reconstructed with UWHAM to obtain *F*(*N*). The HydroMap training label F_dewet_ was taken as the free-energy cost to remove 80% of the hydration-shell waters. The INDUS dataset comprised isolated, capped amino acids (i.e. with ACE and NME caps) in bulk water and 40 protein-embedded residues from hydrophobin and SARS-CoV-2 receptor-binding-domain structures; see Appendix S1.

### HydroMap Model and Training

HydroMap was developed to predict residue-level F_dewet_ from short explicit-solvent molecular dynamics simulations using a small set of interpretable features. The model was trained against INDUS-derived F_dewet_ labels measured for amino acids in two settings: isolated, capped amino acids in bulk water and amino acids embedded in structured proteins. The isolated-residue dataset comprised the 20 canonical amino acids in a99SB-disp and an expanded set across three force fields (a99SB-disp, a03ws, and CHARMM36m), while the structurally diverse dataset comprised 40 protein-embedded residues from hydrophobin and SARS-CoV-2 receptor-binding-domain structures.

For each labeled residue, predictor variables were measured from 5 ns MD simulations. These explicit-solvent simulations were carried out in OpenMM under standard periodic NPT conditions, and the resulting trajectories were used to compute residue–water interaction energies and local water-structure descriptors. Guided by a simple thermodynamic picture in which F_dewet_ reflects a balance between residue–water interaction potential energy and solvent restructuring, HydroMap used sparse linear models with total residue–water potential energy retained as a base feature and one or two additional water-structure descriptors selected from water triplet-angle features by LASSO. Final models were restricted to two or three total features to preserve interpretability and reduce overfitting.

For isolated amino acids in a99SB-disp, the final HydroMap model used total potential energy together with PC1 of the water triplet-angle distribution. To extend this model across force fields, a bulk-water 80–90° triplet-angle feature was added to account for differences in bulk water structuring. For the structurally diverse dataset, the final model used total potential energy together with the 40–50° and 140–150° water triplet-angle fractions. Model performance was evaluated against INDUS F_dewet_ using correlation and root-mean-square error metrics. Models retained their performance with shorter MD simulations. Full details of feature construction, feature selection, dataset composition, and final model comparisons are provided in Appendix S1.

See github.com/samlobe/HydroMap for the open-source model implementation.

### FastHydroMap Model and Training

FastHydroMap was trained to predict HydroMap-derived residue-level F_dewet_ values directly from protein structure, without explicit-solvent simulation at inference time. HydroMap labels were generated for 931 curated PDB proteins, yielding 97,562 residue labels in total; 66,827 residues were retained for supervised modeling and evaluation using protein-level train, validation, and test splits to avoid leakage between residues of the same protein. Retained residues had more than7.0 hydration waters on average, and had F_dewet_ between 3.8 and 8.7 kJ mol^-1^ per water, the approximate range of HydroMap’s training data. These HydroMap labels were generated from 500 ps explicit-solvent simulations per protein.

Each protein was represented as a residue graph with residues as nodes and edges connecting each residue to its 12 nearest neighbors based on Cα distances. Node features encoded amino-acid identity, heavy-atom solvent-accessible surface area (SASA) features, relative SASA features, and terminal flags, while edge features encoded inter-residue distances and relative backbone orientations. The final production model was a compact residue-level message-passing neural network with 46,474 trainable parameters. It predicted F_dewet_ as the sum of an intrinsic term computed from residue-local features before message passing and a contextual term computed after two rounds of message passing.

Training minimized mean-squared error on trusted residues using AdamW with early stopping based on validation RMSE. Model selection and ablations were performed on the validation set only, and the held-out test set was used only after the architecture and hyperparameters had been fixed. Final production weights were then obtained by retraining the locked architecture on the combined training and validation proteins. Performance was summarized using RMSE, Pearson correlation, and Spearman correlation relative to HydroMap labels. For PC1–PC3 analyses, separate single-output regressors were trained using the same architecture, train/validation/test split, trusted-residue mask, optimizer, and early-stopping procedure; PC targets were winsorized to the 2nd–98th percentiles of the trusted training split to reduce the influence of rare extreme values. Full architectural details, hyperparameters, ablations, and training-data fraction analyses are provided in Appendix S2.

See github.com/samlobe/FastHydroMap for the open-source model implementation.

### Case-study structures and trajectories

HydroMap and FastHydroMap were applied to selected static structures and molecular dynamics trajectories to illustrate context-dependent hydrophobicity in biological systems. Static case studies included SARS-CoV-2 receptor-binding-domain structures across major variants, the Parkinson’s disease α-synuclein filament cryo-EM structure (PDB #8A9L), and apo and Ca^2+^-bound calmodulin structures (PDBs #1CFD and #1CLL). Dynamic analysis was performed on a Protein G GB1 folding trajectory from D. E. Shaw Research (49). Per-residue F_dewet_ values were evaluated directly from structure using FastHydroMap or from short explicit-solvent simulations using HydroMap, as described above.

## Acknowledgments

We acknowledge support from the National Institute of Health Grant 5R01AG056058-09, the NSF Grant NSF-ANR MCB/PHY 2423885, the MRSEC Program of the NSF under Award No. DMR 2308708 (IRG-2), and the generous support of a private donor. S.L. acknowledges support from the University of California Graduate Opportunity Fellowship and the Connie Frank Fellowship. J.-E.S. acknowledges support from the NSF (MCB-1716956). M.S.S. gratefully acknowledges funding support from the NSF through award no. CHEM-1800344. We acknowledge the computational facilities purchased with funds from the NSF (CNS-1725797) and administered by the Center for Scientific Computing (CSC). The CSC is supported by the California NanoSystems Institute and the Materials Research Science and Engineering Center (MRSEC; NSF DMR 2308708) at UC Santa Barbara. This work used Anvil at Purdue University and Stampede2 at the Texas Advanced Computing Center through allocation MCA05S027 from the Advanced Cyberinfrastructure Coordination Ecosystem: Services & Support (ACCESS) program, which is supported by U.S. National Science Foundation grants #2138259, #2138286, #2138307, #2137603, and #2138296.

