## Supplemental Information for "Context-Aware Hydrophobicity Modeling: HydroMap and FastHydroMap"

### Supporting Information for
*Context-Aware Hydrophobicity Modeling: HydroMap and FastHydroMap*

Samuel Lobo, Saeed Najafi, Joan-Emma Shea, and M. Scott Shell

Corresponding Authors:
Joan-Emma Shea


M. Scott Shell


**This PDF file includes:**

Figures S1 to S17
Tables S1 to S8
Appendix S1: HydroMap model training (MD-based)
Appendix S2: FastHydroMap model training (MPNN-based)
SI References

**Other supporting materials for this manuscript include the following:**

Movie S1
Software S1 to S2

**Figures**

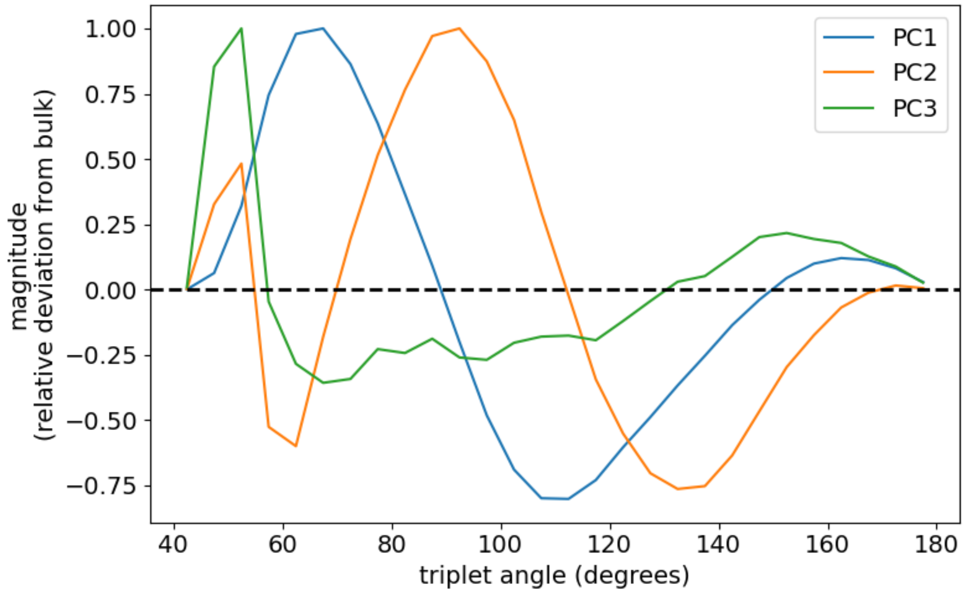


**Fig S1.** The principal components of the water triplet distribution were derived by observing the modes by which water structures upon changes in temperature and density (1). PC1 quantifies the tradeoff between icosahedral waters (~60°) and tetrahedral waters (~109.5°). PC2 is sensitive to 90° waters (prominent in ideal gases and extended surfaces with dangling hydrogen bonds) which tradeoff against icosahedral waters and waters with larger angles. PC3 is sensitive to 50° angles; these associated with a 5$^{th}$ coordinated water and hydrophilic residues.

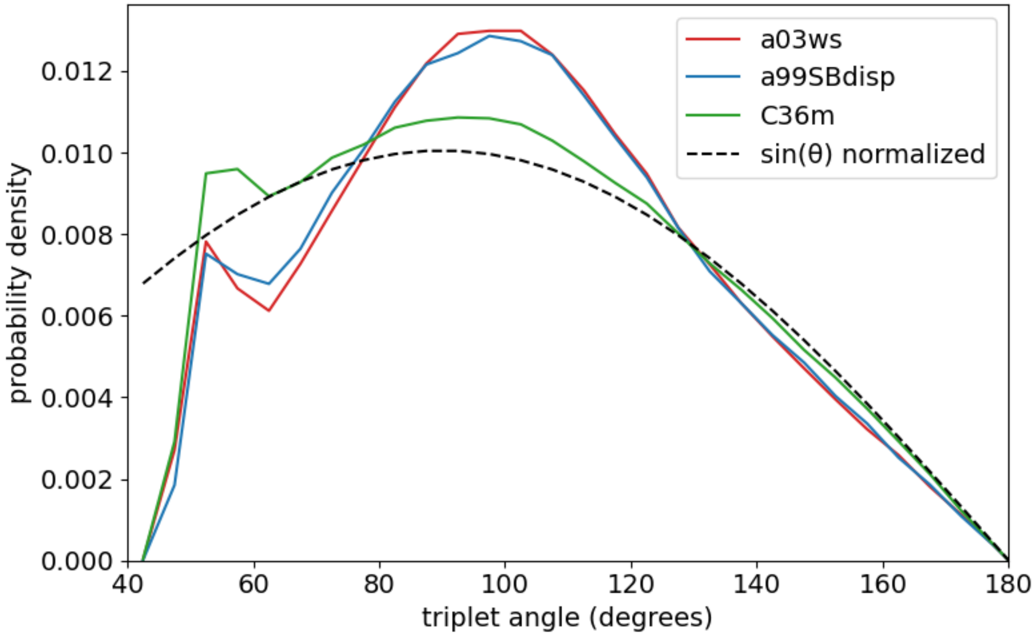


**Fig S2.** The bulk water triplet angle distribution for the a03ws, a99SB-disp, and CHARMM36m force fields. Ideal gases have a triplet distribution according to sin(θ) and it is shown for comparison, normalized to have unit area between 40-180°.

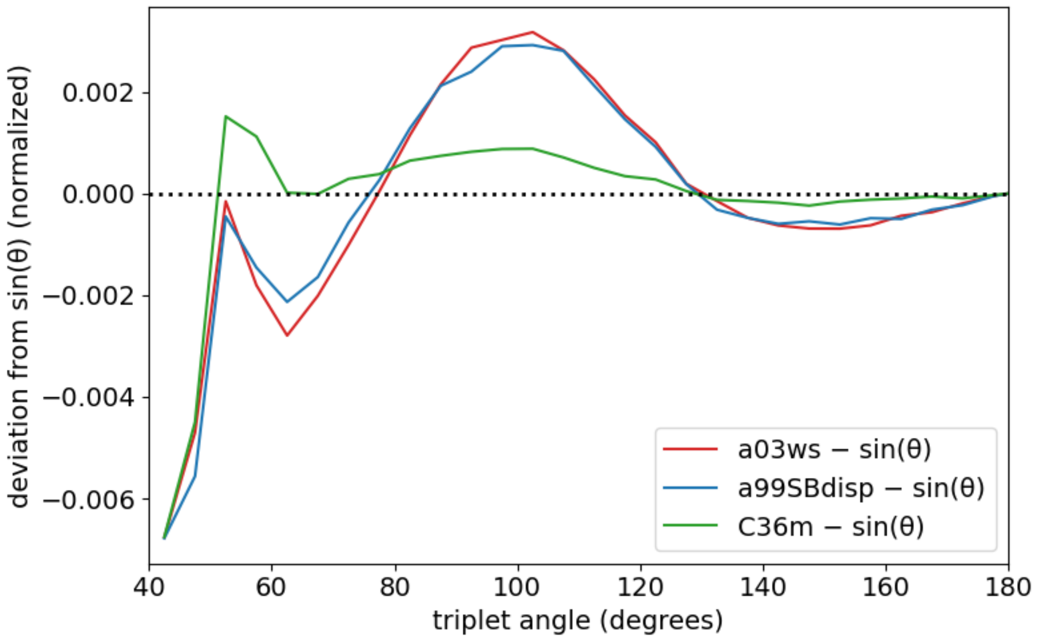


**Fig S3.** Comparing the deviation of the a03ws, a99SB-disp, and CHARMM36m force fields relative an ideal gas distribution (i.e. a normalized sin(θ)). CHARMM36m's bulk water, consisting of TIP3P, has the strongest resemblance to the ideal gas triplet distribution. Higher frequencies of 80-90° bulk water angles are correlated to higher deviations from the ideal gas distribution, indicating more structured, lower-entropy bulk water.

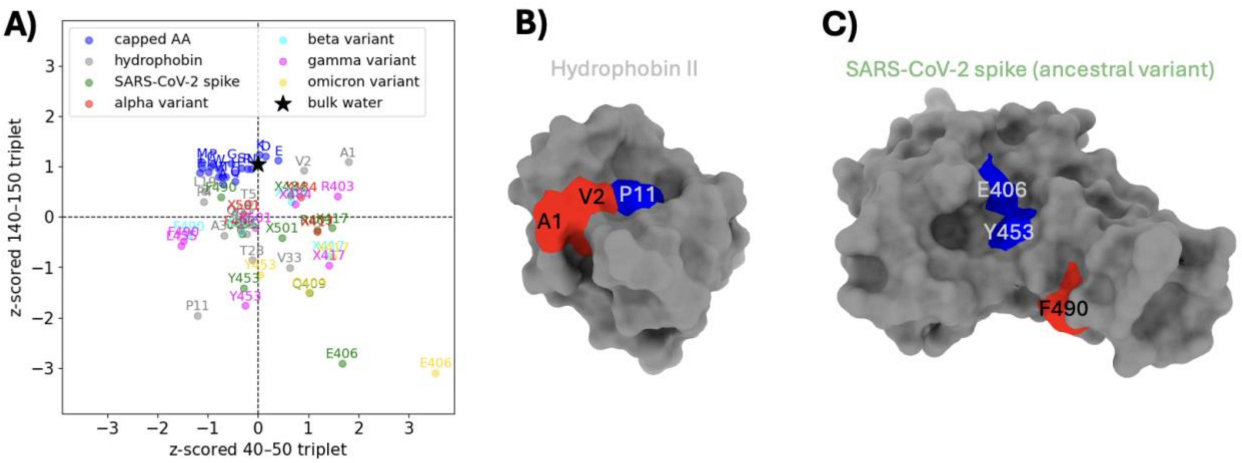


**Fig S4.** A) The z-scored features (140-150° and 40-50° triplet angle fractions) of each amino acid in the HydroMap training data along with bulk water. The isolated, capped amino acids have 140-150° signatures similar to bulk water; there is less depletion of 140-150° angles in small hydrophobes. For context, the mean 40-50° signature and 140-150° signatures in the training set were 0.94% and 4.8% of all the triplet angles, respectively; their standard deviations were 0.12% and 0.37%, respectively. B) Highlighting three residues (A1, V2, P11) in the protein hydrophobin: A1 & V2 have a bulk-like 140-150° signature, while P11 is more depleted in 140-150° water angles. C) Highlighting three residues (E406, Y453, F490) in the SARS-CoV-2 RBD protein (ancestral variant): F490 has a 140-150° signature that is somewhat similar to bulk water, while Y453 & especially E406 are significantly more depleted in 140-150° water angles. It appears that the more bulk-like residues (highlighted red in B and C) are part of more convex surfaces while the 140-150°-depleted residues (highlighted blue) are part of flatter, more concave surfaces.

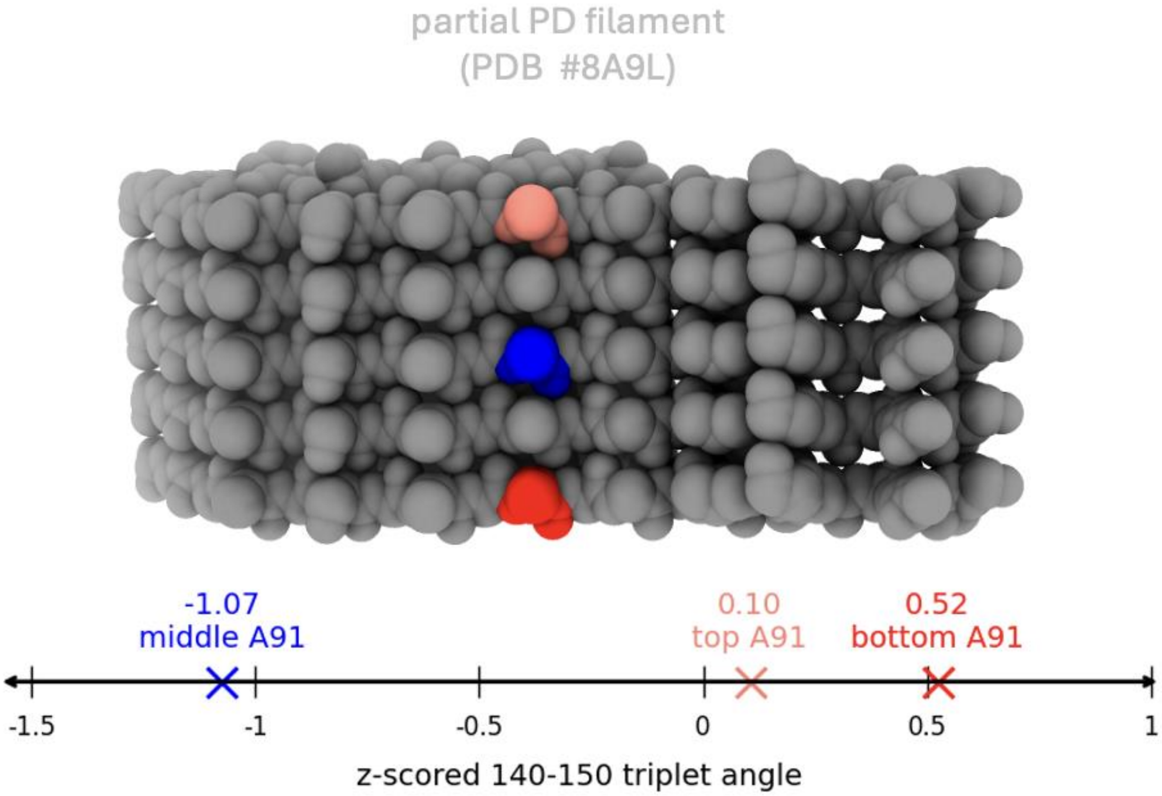


**Fig S5.** Evaluating the 140-150° water signal around an amyloid filament: the α-synuclein filament resolved by cryo-EM from Parkinson's disease patients (PDB #8A9L). This is a partial filament from residues 73 to 100. The middle A91 residue is in a flat, somewhat concave surface and is the most depleted in 140-150° water triplet angles; meanwhile the outer chains' A91 have more bulk-like 140-150° signatures; especially the bottom chain's which is the most convex of the 5 chains.

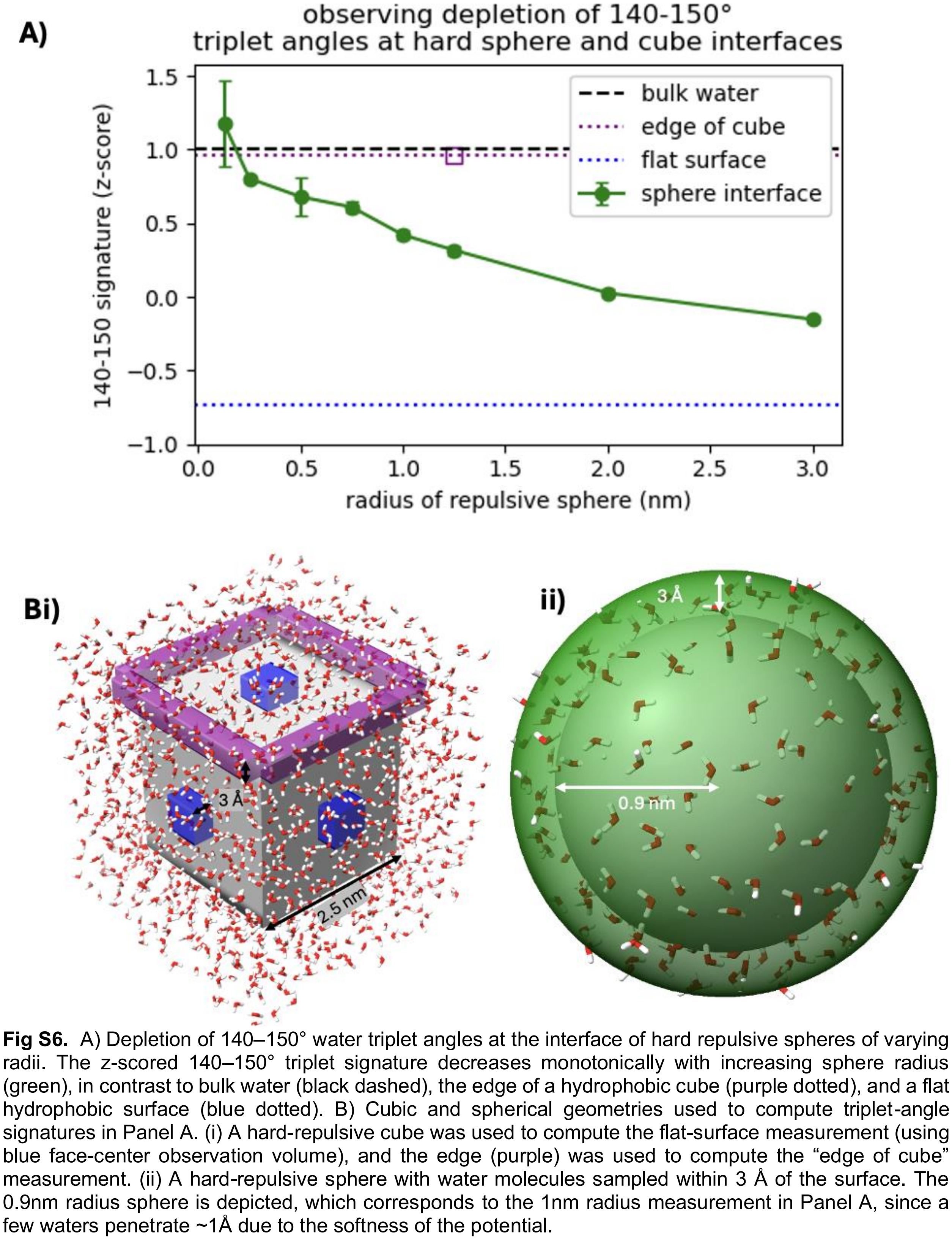


**Fig S6.** A) Depletion of 140–150° water triplet angles at the interface of hard repulsive spheres of varying radii. The z-scored 140–150° triplet signature decreases monotonically with increasing sphere radius (green), in contrast to bulk water (black dashed), the edge of a hydrophobic cube (purple dotted), and a flat hydrophobic surface (blue dotted). B) Cubic and spherical geometries used to compute triplet-angle signatures in Panel A. (i) A hard-repulsive cube was used to compute the flat-surface measurement (using blue face-center observation volume), and the edge (purple) was used to compute the "edge of cube" measurement. (ii) A hard-repulsive sphere with water molecules sampled within 3 Å of the surface. The 0.9nm radius sphere is depicted, which corresponds to the 1nm radius measurement in Panel A, since a few waters penetrate ~1Å due to the softness of the potential.

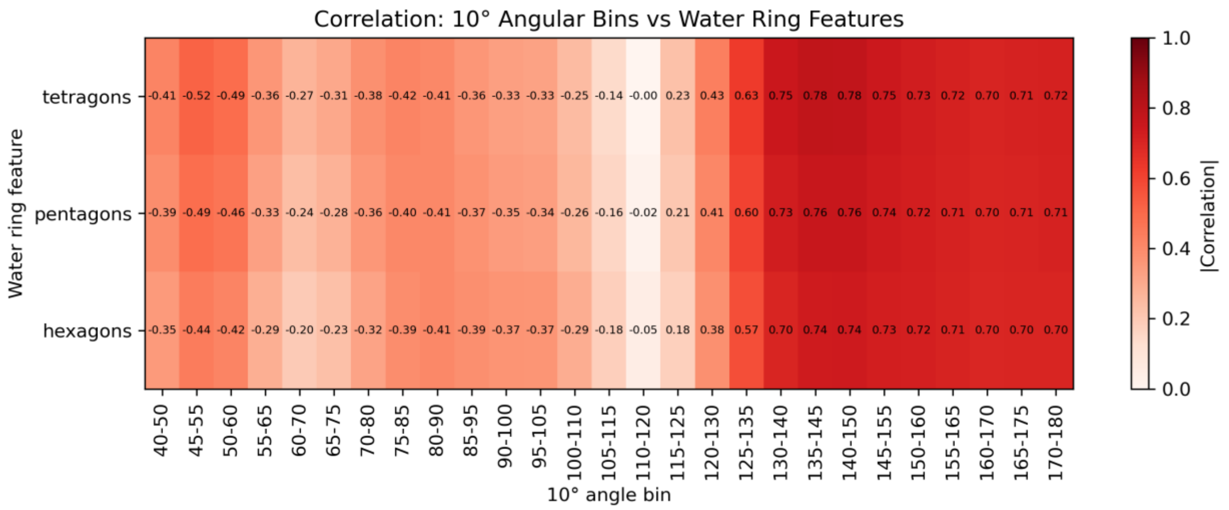


**Fig S7.** Heatmap of correlations between 10° bins of water's triplet distribution and water ring features. 140-150° has the strongest correlations to tetragonal, pentagonal, and hexagonal rings, respectively.

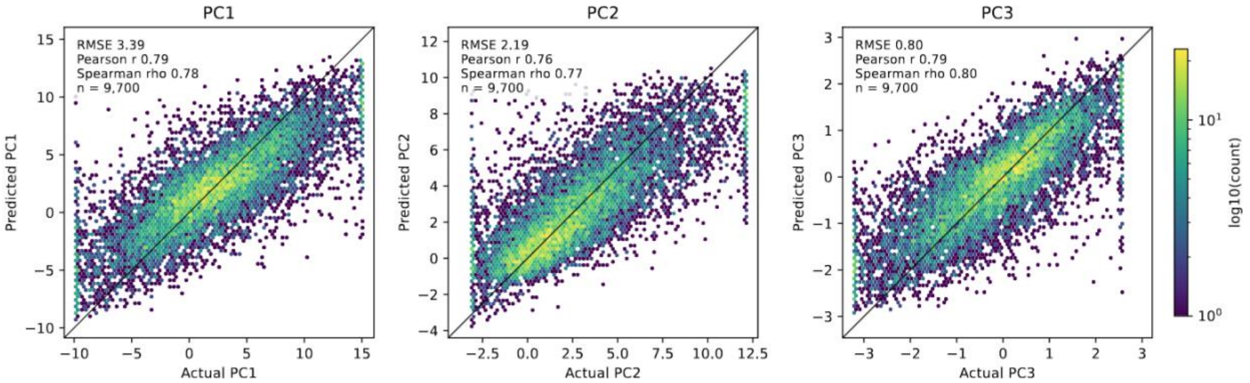


**Fig S8.** FastHydroMap prediction of water triplet-angle principal components. Held-out test-set predictions of HydroMap-derived PC1, PC2, and PC3 using the same FastHydroMap message-passing architecture used for $F_{dewet}$ prediction. Each trusted residues' PC values were regressed, and hexbin color indicates the log10 number of residues in each bin. PC targets were winsorized to the 2nd–98th percentiles of the trusted training split before training to reduce the influence of rare extreme values. Test-set performance was PC1: RMSE = 3.39, Pearson r = 0.79, Spearman ρ = 0.78; PC2: RMSE = 2.19, Pearson r = 0.76, Spearman ρ = 0.77; and PC3: RMSE = 0.80, Pearson r = 0.79, Spearman ρ = 0.80.

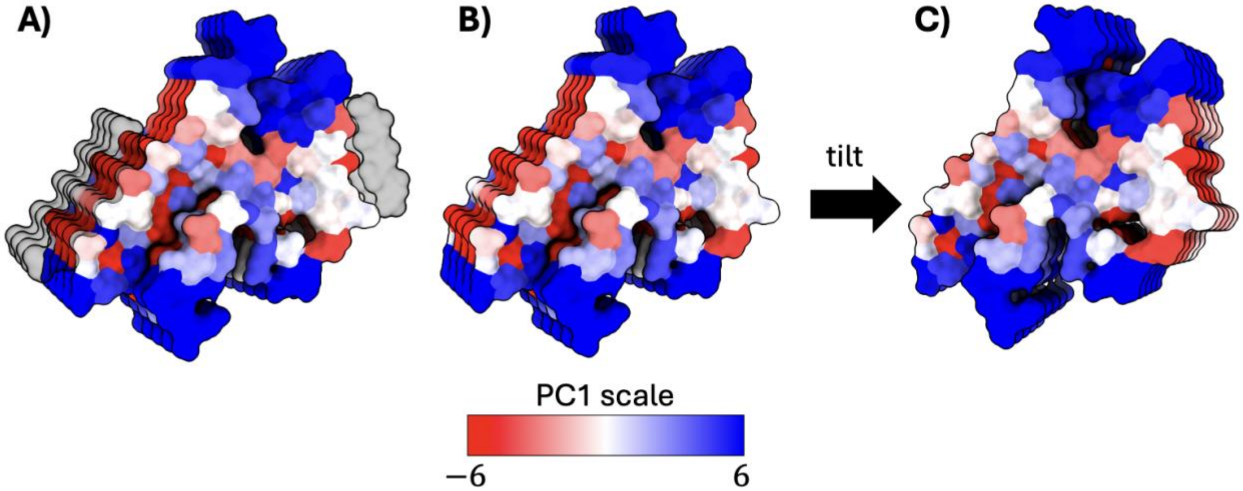


**Fig S9.** There are two interfaces on the α-synuclein Parkinson's amyloid structure (PDB #8A9L) that make contact with unassigned peptide densities. At these interfaces, water structure is significantly more tetrahedral, as indicated by negative PC1 values (see Fix S1). The α-synuclein cryo-EM structure (PDB #8A9L) with (A) and without (B) the unassigned densities is shown along with a tilted version (C).

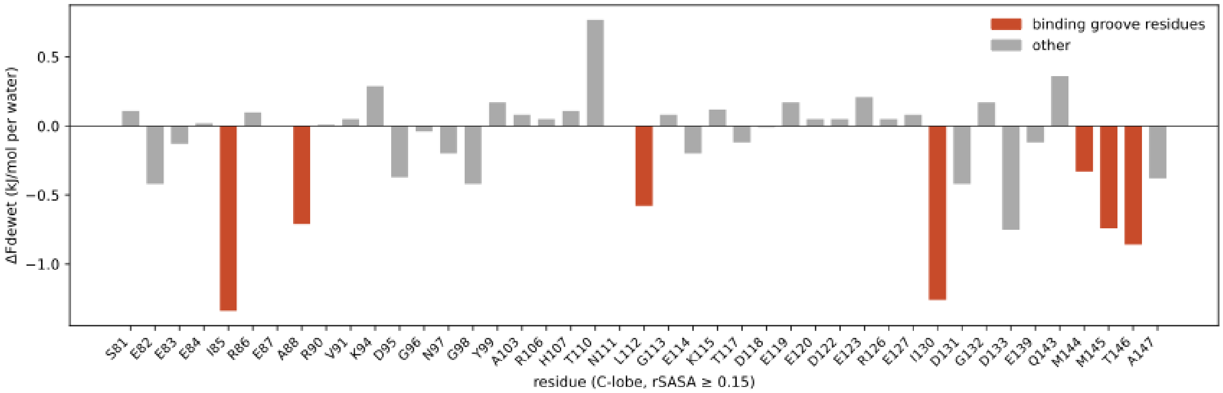


**Fig S10.** $\Delta F_{dewet}$ upon $Ca^{2+}$ binding across C-lobe residues of calmodulin (residues 81–147) with open-state relative solvent-accessible surface area (rSASA) ≥ 0.15. Each bar shows the change in FastHydroMap-predicted dewetting free energy between the apo and $Ca^{2+}$-bound crystal structures ($\Delta F_{dewet} = F_{dewet,holo} - F_{dewet,apo}$). Residues comprising the methionine-rich helix-binding groove are highlighted in red; these residues become more hydrophobic upon $Ca^{2+}$ binding and the resulting conformation change.

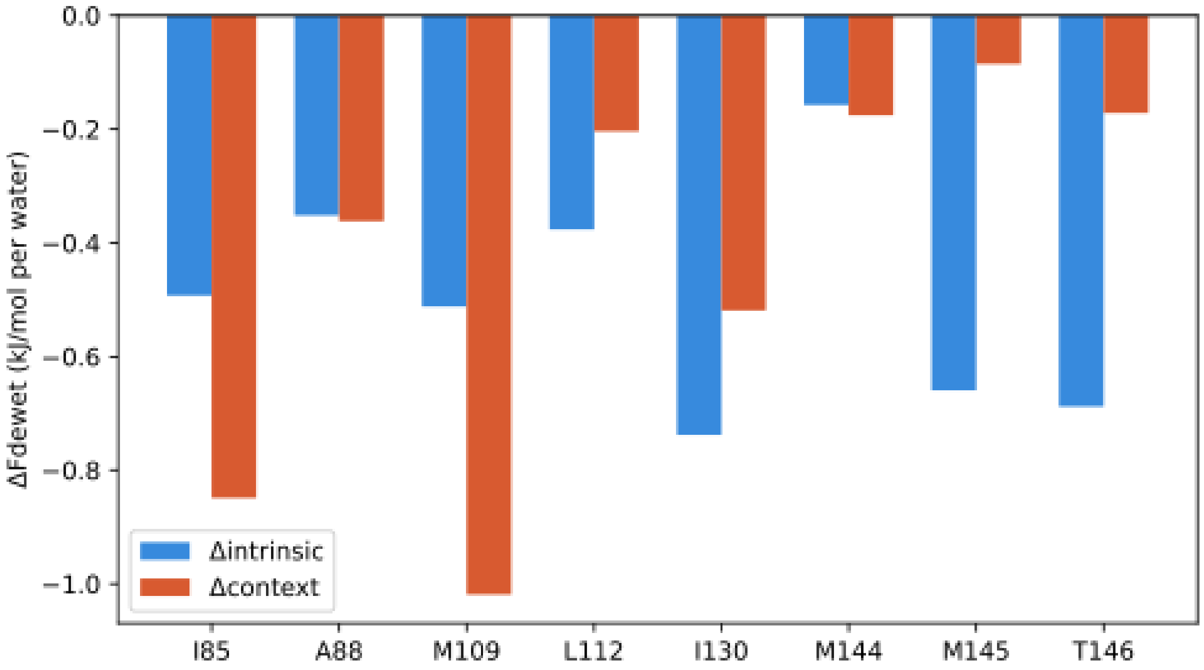


**Fig S11.** Decomposition of $\Delta F_{dewet}$ into intrinsic and context contributions for binding groove residues highlighted in calmodulin's C-lobe. For each highlighted residue, $\Delta F_{dewet}$ is decomposed into a change in the intrinsic term ($\hat{F}_{intrinsic}$; blue), reflecting baseline dewetting propensity from amino acid identity and solvent exposure, and a change in the context term ($\hat{F}_{context}$; orange), reflecting neighbor-dependent modulation after message passing in FastHydroMap. Residues such as M109 and I85 are predominantly context-driven, reflecting changes in neighboring residues, while M145 and T146 are predominantly intrinsic-driven, reflecting changes in their solvent exposure. Negative $\Delta F_{dewet}$ values correspond to increased hydrophobicity in the $Ca^{2+}$-bound conformation.

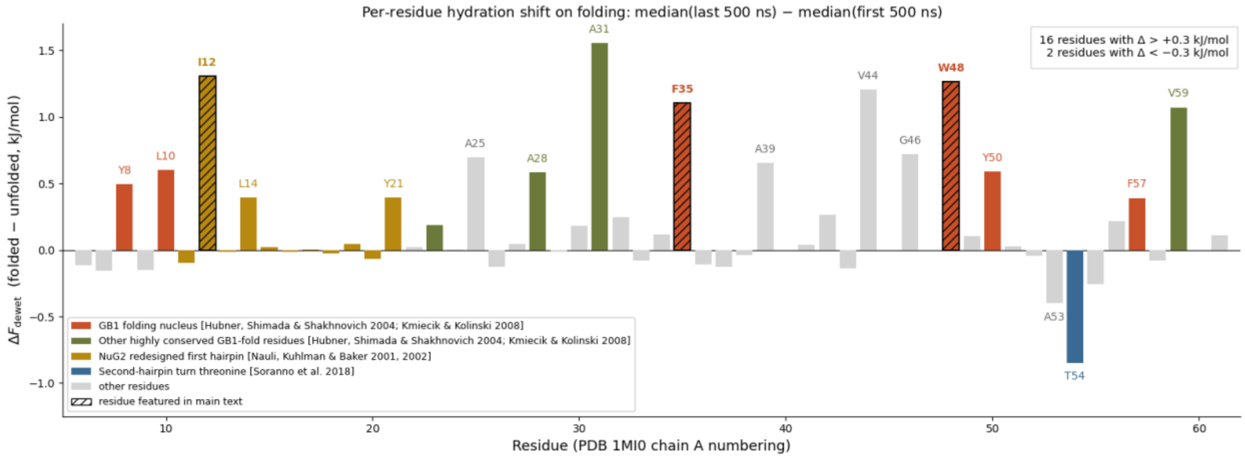


**Fig S12.** Per-residue change in $F_{dewet}$ during the Protein G GB1 domain folding trajectory, quantified as the median value in the last 500 ns minus the median value in the first 500 ns. Residue numbering follows PDB #1MI0. Hatched bars mark the three residues highlighted in Figure 6 of the main text (I12, F35, W48). Colored bars indicate residues with literature-supported structural roles: the conserved GB1 folding nucleus retained in NuG2 (red; Y8, L10, F35, W48, Y50, F57 in this PDB numbering, corresponding to canonical Y3, L5, F30, W43, Y45, F52), additional highly conserved residues of the GB1 fold (green; T23, A28, A31,

V59, corresponding to canonical T18, A23, A26, V54), the 11-residue NuG2 redesigned first hairpin (gold; residues 11–21), and the second-hairpin turn threonine (blue; T54, corresponding to canonical T49) (2–5). Most large $F_{dewet}$ shifts are positive, consistent with hydrophobic surfaces being buried away from solvent while more water-compatible surfaces remain exposed upon folding. Many of the large upward shifts fall in residues identified by prior simulation and sequence-conservation analyses as the GB1 folding nucleus and its associated conserved core (Y8, L10, F35, W48, Y50, A28, A31, V59) (2, 3). A small number of residues shift in the opposite direction; T54 shows the largest such shift.

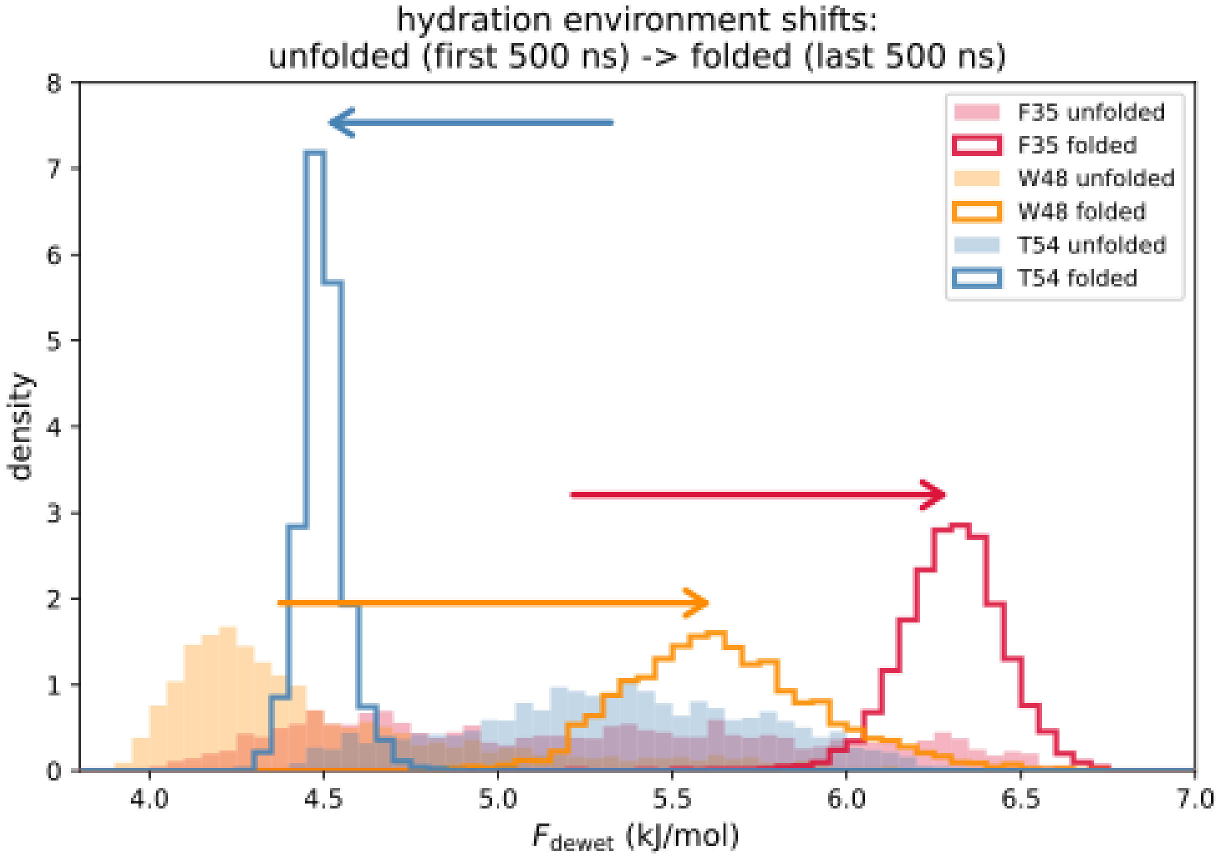


**Fig S13.** Distributions of $F_{dewet}$ in the unfolded (first 500 ns, filled) and folded (last 500 ns, outlined) segments of the Protein G folding trajectory, for three illustrative residues. F35 (red) and W48 (orange) both shift to higher $F_{dewet}$ upon folding, consistent with their hydrophobic side chains burying and thereby displaying less hydrophobic surfaces to water. Both distributions also narrow upon folding, reflecting changes in local environments, though W48's folded distribution remains broader than F35's. T54 (blue) shifts in the opposite direction and sharpens dramatically, consistent with threonine being locking into a specific, solvent-exposed geometry. Arrows indicate the direction of each distribution shift during folding.

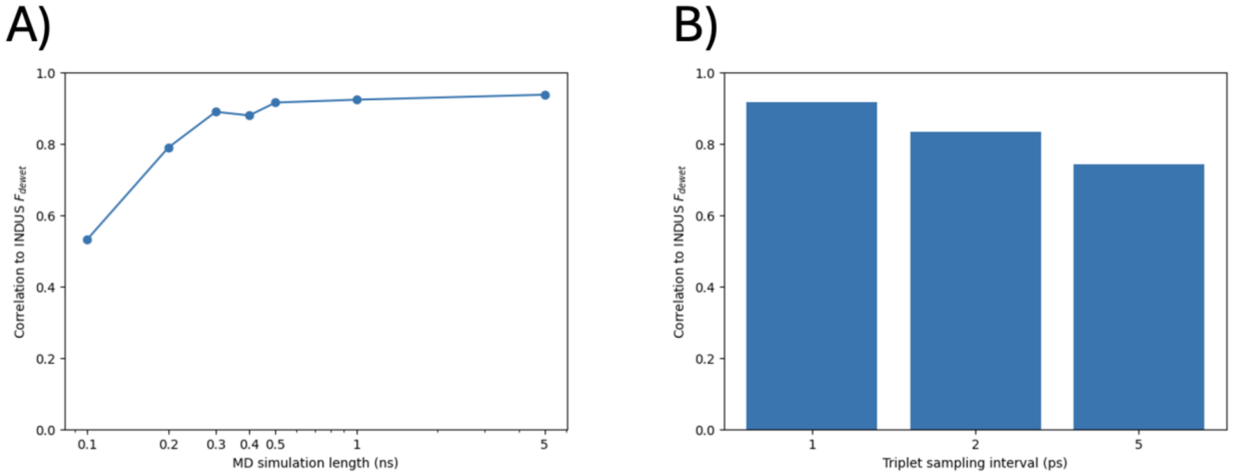


**Fig S14.** Sampling requirements for accurate HydroMap predictions of residue-level dewetting free energy. A) Correlation between HydroMap-predicted and INDUS-computed $F_{\mathrm{dewet}}$ as a function of MD trajectory length used to compute water-structure features. High accuracy is achieved with only a few hundred picoseconds of sampling, with performance saturating by ~0.5–1 ns. B) Sensitivity of prediction accuracy to temporal subsampling of water configurations. Correlation remains high even when triplet-angle statistics are computed at coarse sampling intervals (1–5 ps), indicating that detailed, high-frequency sampling of water structure is not required.

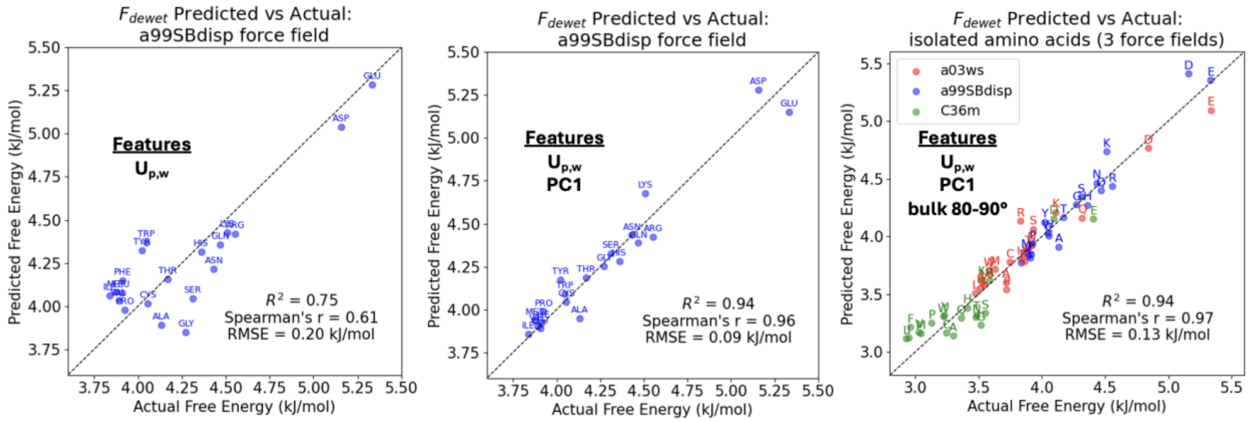


**Fig S15.** HydroMap performance on isolated amino acids across feature sets and force fields. $F_{dewet}$ predicted by HydroMap versus reference INDUS values for each of the 20 standard amino acids in their isolated tripeptide form. Left) Performance using only the protein–water interaction energy U_p,w as input feature (a99SBdisp force field), $R^2 = 0.75$, Spearman's r = 0.61, RMSE = 0.20 kJ/mol per water. Middle) Adding the first principal component of the local water-structure features (PC1) substantially improves prediction (a99SBdisp), $R^2 = 0.94$, Spearman's r = 0.96, RMSE = 0.09 kJ/mol per water. Right) Including the bulk 80–90° triplet angle population further generalizes the model across three commonly used force fields (a03ws, a99SBdisp, C36m), $R^2 = 0.94$, Spearman's r = 0.97, RMSE = 0.13 kJ/mol per water. Dashed line indicates y = x. Each point corresponds to one amino acid (one-letter code).

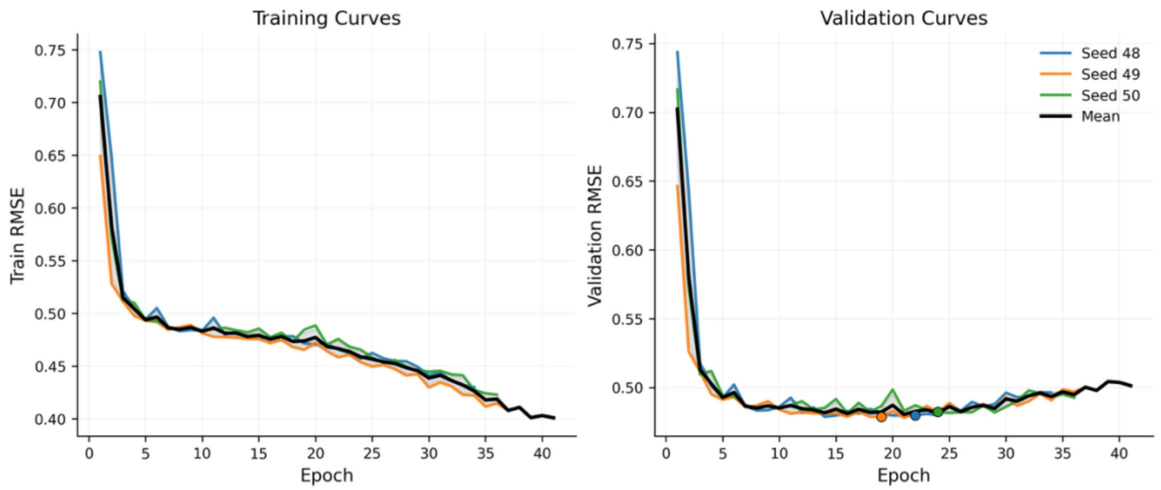


**Fig S16.** Validation-set loss curves across three random initializations for the locked FastHydroMap architecture. Curves should show validation RMSE versus epoch for seeds 48, 49, and 50, with best-epoch markers indicated.

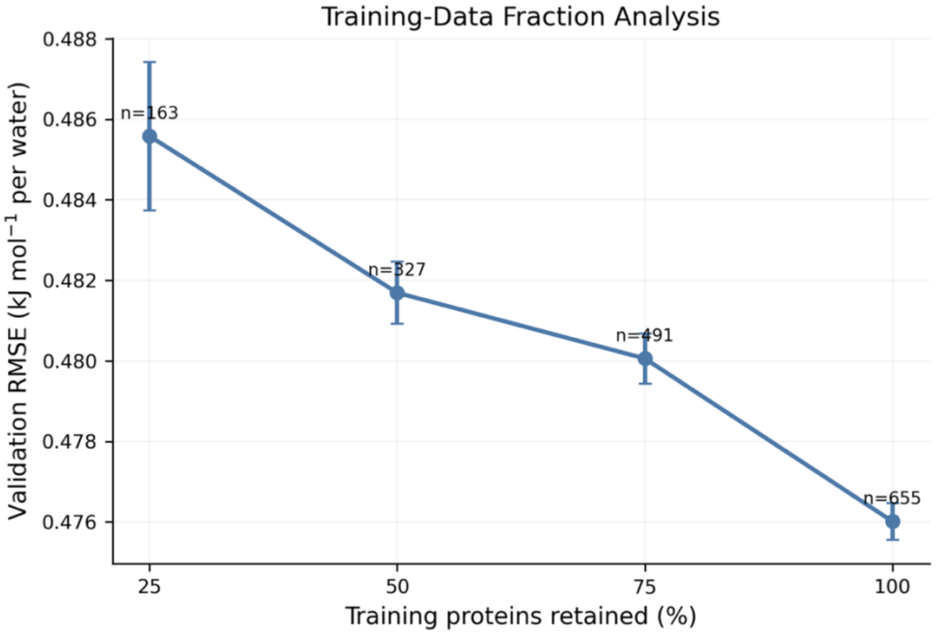


**Fig S17.** Training-data fraction analysis for FastHydroMap. Validation RMSE as a function of the fraction of training proteins retained (25%, 50%, 75%, 100%), with error bars indicating standard deviation across three seeds.

**Appendix S1:** HydroMap model training (MD-based)

##### S1.1 HydroMap training datasets and simulation-derived features

HydroMap was trained against INDUS-derived residue-level $F_{\mathrm{dewet}}$labels measured in two settings: isolated, capped amino acids in bulk water and amino acids embedded in structured proteins. The isolated-residue dataset comprised the 20 canonical amino acids simulated in a99SB-disp, and an expanded set across three force fields (a99SB-disp, a03ws, and CHARMM36m), yielding 60 total isolated-residue conditions. The structurally diverse dataset comprised 40 protein-embedded residues from hydrophobin and SARS-CoV-2 receptor-binding-domain (RBD) structures across multiple variants; the full residue list is given in Table S1. All protein-embedded residues were simulated in a99SB-disp.

For each labeled residue, predictor variables were measured from short, unbiased explicit-solvent MD simulations. Independent 5 ns simulations were used for the final feature collection. Final HydroMap feature-collection simulations were performed in OpenMM using explicit solvent and periodic boundary conditions. Simulations were run in the NPT ensemble at 300 K and 1 atm with PME electrostatics and a standard nonbonded cutoff. Protein-embedded systems were simulated with restraints to preserve the local geometry used in the corresponding INDUS label generation. Potential energy terms were measured from these trajectories, and water-structure fingerprints were computed within approximately two hydration layers of each residue.

Potential energy terms were measured with OpenMM, and water structural fingerprints were measured within approximately two hydration layers of the residue. The final HydroMap fitting used linear regression models with a small number of interpretable features.

| **source** | **amino acids** | **structural context** |
|---|---|---|
| isolated, capped amino acids | A, C, D, E, F, G, H, I, K, L, M, N, P, Q, R, S, T, V, W, Y | isolated |
| Hydrophobin | A1, V2, P4, T5, P11, L19, T28, A32, V33, A37, Q40, Q65 | protein-embedded |
| SARS-CoV-2 RBD (ancestral) | R403, E406, Q409, K417, Y453, L455, E484, F490, N501 | protein-embedded |
| SARS-CoV-2 RBD (Alpha) | K417, E484, F490, Y501 | protein-embedded |
| SARS-CoV-2 RBD (Beta) | N417, K484, F490, Y501 | protein-embedded |
| SARS-CoV-2 RBD (Gamma) | R403, T417, Y453, L455, K484, F490, Y501 | protein-embedded |
| SARS-CoV-2 RBD (Omicron) | E406, Q409, N417, Y453 | protein-embedded |

Table S1. Amino acids in structurally-diverse model set for INDUS $F_{\mathrm{dewet}}$ regression.
The dataset comprises 20 isolated, capped amino acids and 40 protein-embedded amino acids drawn from hydrophobin and SARS-CoV-2 receptor binding domain (RBD) structures across multiple variants. Protein-embedded residues span diverse amino acid chemistries and local structural environments.

##### S1.2 Candidate feature pool and fitting strategy

The final candidate feature pool consisted of residue–water potential energy terms together with water triplet-angle descriptors. Specifically, we considered total, Coulombic, and Lennard-Jones residue–water interaction energies; 27 overlapping 10° bins of the water triplet-angle distribution (e.g. 40-50°, 45-55°, …, 170-180°); and the first three principal components of that distribution taken from Jiao, et al. (6) In earlier exploratory screening, we also considered hydrogen-bond and water-ring descriptors (i.e. tetragonal, pentagonal, and hexagonal water rings in the hydration shell, via github.com/vitroid/CountRings) (7), but these were not retained in the reduced final candidate set used for model training.

Feature selection was guided by both statistical performance and physical interpretability. Total potential energy was always retained as a base feature because it corresponds naturally to the residue–water interaction term in the thermodynamic picture motivating HydroMap. Additional water-structure features were then selected with LASSO to approximate the solvent restructuring contribution. When two candidate features gave nearly identical performance, we favored the more physically interpretable one. To minimize overfitting and preserve interpretability, final HydroMap models were restricted to two or three total features.

##### S1.3 Final model for isolated amino acids in a99SB-disp

For the 20 isolated, capped amino acids simulated in a99SB-disp, a two-feature model consisting of total potential energy and PC1 of the water triplet-angle distribution recovered INDUS $F_{dewet}$ with high accuracy ($R^2$ = 0.94; RMSE = 0.09 kJ $mol^{-1}$ per water). Total potential energy alone achieved $R^2$ = 0.75 under leave-one-out cross-validation, and the best-performing second feature was the 65–75° triplet-angle bin. We nevertheless selected PC1 instead because it gave nearly identical performance ($R^2$ = 0.94) while remaining more physically interpretable.

PC1 captures a tradeoff between more icosahedral, simple-fluid-like hydration environments and more tetrahedral, water-like environments. In this dataset, more negative PC1 values correspond to more tetrahedral local water structure and lower $F_{\mathrm{dewet}}$, whereas more positive PC1 values correspond to more icosahedral local structure and higher $F_{\mathrm{dewet}}$. Thus, for small hydrophobes, tetrahedral ordering around the solute is associated with greater dewetting propensity.

##### S1.4 Final model for isolated amino acids across three force fields

To extend the isolated-residue HydroMap model across force fields with different bulk-water structure, one additional bulk-water descriptor was required. Among bulk triplet-angle features, the bulk 80–90° fraction provided the best correction, yielding a three-feature model with $R^2$ = 0.94 across the 60 isolated amino acid conditions from a99SB-disp, a03ws, and CHARMM36m.

We interpret this bulk 80–90° term as correcting for force-field-dependent differences in bulk water configurational entropy. CHARMM36m, which uses TIP3P water, exhibits a flatter bulk triplet-angle distribution closer to the ideal-gas $\sin(\theta)$ form, whereas a99SB-disp and a03ws show stronger structuring over approximately 80–125°. A larger bulk 80–90° fraction therefore reflects more structured, lower-entropy bulk water, which reduces the entropy contrast between the hydration shell and the bulk and opposes dewetting. In this way, the bulk 80–90° feature captures systematic shifts in hydrophobic driving force across force fields.

##### S1.5 Final model for structurally diverse residues

For the structurally diverse dataset comprising protein-embedded residues together with isolated amino acids, the final HydroMap model used three features: total potential energy, the 40–50° triplet-angle fraction, and the 140–150° triplet-angle fraction. LASSO repeatedly selected an potential energy term together with one low-angle and one high-angle triplet feature from this dataset; in the final reduced candidate pool, the selected features were Coulombic potential energy, 40–50°, and 140–150°. For simplicity and physical consistency across HydroMap models, we replaced Coulombic potential energy with total potential energy with negligible loss of performance. This final three-feature model achieved $R^2$ = 0.88, and performance remained essentially unchanged when the isolated amino acids were included jointly with the protein-embedded residues.

The 40–50° feature is associated with highly coordinated hydration waters and is strongest at polar and charged sites, consistent with prior interpretations of ~48° triplet angles as signatures of hyper-coordinated water (8). In our dataset, larger 40–50° populations are associated with higher $F_{dewet}$ and thus with hydrophilicity. The 140–150° feature behaves oppositely: it is strongest in bulk water and around small hydrophobes, but is depleted in flatter or more concave protein environments. Residues such as A1 and V2 in hydrophobin and solvent-exposed F490 in the SARS-CoV-2 RBD retain relatively bulk-like 140–150° populations, whereas residues such as P11 in hydrophobin and E406 in the SARS-CoV-2 RBD show strong depletion of this feature. Across the full dataset, the 140–150° feature is strongly anticorrelated with PC2 and positively correlated with hydrogen-bonded water-ring features, suggesting that it reports on topology-sensitive restructuring of interfacial water.

Taken together, these HydroMap models show that residue–water interaction energy plus one or two water-structure fingerprints suffice to recover INDUS $F_{dewet}$ across isolated and protein-embedded contexts. The resulting models remain sparse and interpretable while capturing both chemical and geometric contributions to context-dependent hydrophobicity.

Table S2. Summary of final HydroMap models

| dataset | residues | final features | performance | notes |
|---|---|---|---|---|
| Isolated amino acids (a99SB-disp) | 20 | $U_{pw}$, PC1 | $R^2 = 0.94$; RMSE = 0.09 kJ mol$^{-1}$ per water | PC1 was chosen over the slightly better-performing 65–75° bin because it gave nearly identical accuracy while remaining more physically interpretable. |
| isolated amino acids (3 force fields) | 60 | $U_{pw}$, PC1, bulk 80–90° | $R^2 = 0.94$; RMSE = 0.13 kJ mol$^{-1}$ per water | The bulk 80–90° term corrects for force-field-dependent differences in bulk water structuring and entropy. |
| structurally diverse residues (protein-embedded only) | 40 | $U_{pw}$, 40–50° triplet, 140–150° triplet | $R^2 = 0.88$ | Final reduced-candidate-set fit for protein-embedded residues from hydrophobin and SARS-CoV-2 RBD variants. |
| structurally diverse residues + isolated amino acids | 60 | $U_{pw}$, 40–50° triplet, 140–150° triplet | $R^2 = 0.88$; Spearman r = 0.89; RMSE = 0.33 kJ mol$^{-1}$ per water | Performance was essentially unchanged when isolated residues were included jointly with the protein-embedded set. |

**Appendix S2:** FastHydroMap model training (MPNN-based)

#### S2.1 Dataset construction and filtering

HydroMap labels were generated for 931 curated PDB proteins. Proteins were filtered to include single-chain structures with fewer than 200 amino acids, no missing amino acids, no non-canonical amino acids, no additional organic molecules, and less than 50% pairwise sequence similarity. Across these proteins, HydroMap produced 97,562 residue-level labels in total.

For supervised modeling and evaluation, we retained a trusted subset of residues within the $F_{\text{dewet}}$and solvation range of HydroMap's training data. A residue was considered trusted if its average hydration-water count exceeded 7.0 and its HydroMap $F_{\text{dewet}}$ value fell between 3.8 and 8.7 kJ $\text{mol}^{-1}$ per water. This yielded 66,827 trusted residues.

Protein-level train, validation, and test splits were used throughout to avoid leakage between residues from the same protein. The split contained 655 training proteins, 141 validation proteins, and 141 test proteins. All model selection, hyperparameter tuning, and ablation comparisons were performed on the validation set only. The held-out test set was used only after the architecture and hyperparameters had been fixed.

#### S2.2 Graph features and architecture

Each protein was represented as a residue graph in which each residue was a node. Edges connected each residue to its 12 nearest neighbors in C$\alpha$ space. Graph construction was chain-aware and insertion-code-aware, so multichain inputs and residues with insertion codes were handled using unique residue identifiers. Solvent-accessible surface areas were featurized using heavy atoms only.

Each residue was represented by a 32-dimensional node feature vector consisting of: (i) a 20-dimensional one-hot encoding over the canonical amino acids; (ii) raw heavy-atom SASA features for backbone N, C$\alpha$, C, O, and C$\beta$, together with side-chain carbon SASA beyond C$\beta$ ($sc_c$) and side-chain N/O/S SASA ($sc_{NOS}$); (iii) residue-standardized versions of total residue SASA, $sc_c$ SASA, and $sc_{NOS}$ SASA; and (iv) binary N-terminal and C-terminal flags.

Each edge was represented by an 84-dimensional feature vector consisting of: (i) 25 pairwise distances between atoms drawn from $\{\text{N}, \text{C}\alpha, \text{C}\beta, \text{C}, \text{O}\}$ across the residue pair; (ii) a 3-bin radial basis function expansion of each distance, using centers spanning 2–14 Å with $\sigma = 4$Å; and (iii) nine relative orientation features derived from local backbone frames.

The final production model was a residue-level message-passing neural network with 46,474 trainable parameters. Node features were first projected into a hidden embedding space of dimension 24. An intrinsic prediction head operated on this projected embedding before message passing. Two rounds of mean-aggregation message passing were then applied, followed by a contextual prediction head on the final message-passed embedding. The final prediction for each residue was

$$F_{\text{dewet}} = \mu + F_{\text{intrinsic}} + F_{\text{context}},$$

where $\mu$ is the mean trusted $F_{\text{dewet}}$ value in the training set. The hidden dimension was 24, the message-passing depth was 2, and the hidden dimension of the prediction heads was 20.

#### S2.3 Training procedure and final performance

Training minimized mean-squared error on trusted residues only. Optimization used AdamW with weight decay and a Noam-style learning-rate schedule. Gradient clipping, dropout, and edge dropout were applied during training. See Table S3.

Table S3. FastHydroMap hyperparameters

| Parameter | Value |
|---|---|
| Graph neighbors (k) | 12 |
| Node feature dimension | 32 |
| Edge feature dimension | 84 |
| Hidden dimension | 24 |

| Parameter | Value |
|---|---|
| Message-passing depth | 2 |
| Head hidden dimension | 20 |
| Distance basis | 3 RBFs |
| RBF range | 2–14 Å |
| RBF sigma | 4 Å |
| Dropout | 0.1 |
| Edge dropout | 0.1 |
| Weight decay | 1 × 10^-3 |
| Warmup | 300 |
| Batch size | 32 |
| Maximum epochs | 50 |
| Patience | 20 |
| Gradient clip | 2.0 |
| Trainable parameters | 46,474 |

Validation-stage runs used early stopping based on validation RMSE. After architecture and hyperparameter selection, the final production weights were obtained by retraining the locked architecture on the combined training and validation proteins for a fixed number of epochs determined from the validation-stage run. In the final production run, this epoch count was 22.

Across three random initializations, the locked FastHydroMap architecture achieved a held-out test RMSE of $0.4719 \pm 0.0010$ kJ mol$^{-1}$ per water, with Pearson $r = 0.7880 \pm 0.0008$ and Spearman $\rho = 0.7897 \pm 0.0003$.

Table S4. Final held-out test performance across three random initializations

| Metric | Mean | Standard deviation |
|---|---|---|
| RMSE (kJ mol^-1 per water) | 0.4719 | 0.0010 |
| Pearson r | 0.7880 | 0.0008 |
| Spearman rho | 0.7897 | 0.0003 |

To test whether the same architecture could predict water-structure descriptors beyond $F_{dewet}$, we trained separate FastHydroMap regressors for PC1, PC2, and PC3 of the water triplet-angle distribution, measured from HydroMap MD simulations. The PC regressors used the same graph representation, architecture, hyperparameters, trusted-residue mask, and protein-level train/validation/test splits as the Fdewet model. Because a minority of residues had extreme PC values, PC targets were winsorized using the 2nd and 98th percentiles of the trusted training split before training. The resulting clipping bounds were −9.865 to 15.045 for PC1, −3.099 to 12.146 for PC2, and −3.212 to 2.571 for PC3. On the held-out test set, the models achieved Spearman ρ = 0.784, 0.770, and 0.801 for PC1, PC2, and PC3, respectively (Fig. S8).

##### S2.4 Architecture and feature ablations

We evaluated a series of validation-set ablations for the locked FastHydroMap architecture. Core architecture ablations and feature ablations are summarized in Tables S2.3–S2.5. As a label-shuffle control, randomly permuting the training labels degraded validation RMSE to $0.7671 \pm 0.0039$.

Table S5. Core architecture ablations on the validation set

| Model | Parameters | Validation RMSE mean | Validation RMSE s.d. |
|---|---|---|---|
| Full model | 46,474 | 0.4760 | 0.0005 |
| Linear head | 45,482 | 0.4772 | 0.0015 |

| Model | Parameters | Validation RMSE mean | Validation RMSE s.d. |
|---|---|---|---|
| Depth = 1 | 24,154 | 0.4775 | 0.0003 |
| Context only | 45,953 | 0.4782 | 0.0007 |
| Intrinsic only | 1,313 | 0.4811 | 0.0004 |
| Shuffled labels | 46,474 | 0.7671 | 0.0039 |

Table S6. Feature ablations on the validation set

| Model | Node dim | Edge dim | Parameters | Validation RMSE mean | Validation RMSE s.d. |
|---|---|---|---|---|---|
| Full model | 32 | 84 | 46,474 | 0.4760 | 0.0005 |
| No orientation terms | 32 | 75 | 39,382 | 0.4782 | 0.0007 |
| No relative SASA | 29 | 84 | 46,402 | 0.4792 | 0.0010 |
| No terminal flags | 30 | 84 | 46,426 | 0.4799 | 0.0016 |
| Overall SASA only | 24 | 84 | 46,282 | 0.4890 | 0.0002 |

Table S7. Distance encoding ablation on the validation set

| Model | Edge dim | Parameters | Validation RMSE mean | Validation RMSE s.d. |
|---|---|---|---|---|
| 3-bin RBF distances | 84 | 46,474 | 0.4760 | 0.0005 |
| Raw distances | 34 | 15,274 | 0.4810 | 0.0008 |

**S2.5 Training-data fraction analysis**

To assess data efficiency, we repeated validation-stage training with the locked architecture while keeping the validation set fixed and randomly subsampling the training proteins. Reducing the training set from 100% to 75%, 50%, and 25% increased validation RMSE from $0.4760 \pm 0.0005$ to $0.4801 \pm 0.0006$, $0.4817 \pm 0.0008$, and $0.4856 \pm 0.0018$ kJ mol$^{-1}$ per water, respectively.

Table S8. Training-data fraction analysis on the validation set

| Training fraction | Training proteins | Validation proteins | Parameters | Validation RMSE mean | Validation RMSE s.d. |
|---|---|---|---|---|---|
| 100% | 655 | 141 | 46,474 | 0.4760 | 0.0005 |
| 75% | 491 | 141 | 46,474 | 0.4801 | 0.0006 |
| 50% | 327 | 141 | 46,474 | 0.4817 | 0.0008 |
| 25% | 163 | 141 | 46,474 | 0.4856 | 0.0018 |

**Movie S1.** Time-resolved FastHydroMap prediction of residue-level hydrophobicity during Protein G folding. The movie shows the redesigned B1 domain of streptococcal Protein G (GB1) along a partial folding trajectory (~182 ns) from D. E. Shaw Research, colored by FastHydroMap-predicted per-residue dewetting free energy, with yellow representing low dewetting free energies (hydrophobic) and blue representing high dewetting free energies (hydrophilic); bounds of 4.0 to 6.5 kJ/mol per water were used with the lipophilicity palette in ChimeraX. Residues I12, F35, and W48 are highlighted.

**Software S1.** Open-source HydroMap GitHub repository is publicly accessible at https://github.com/samlobo/HydroMap and have been deposited in Zenodo as archival snapshots.

**Software S2.** Open-source FastHydroMap GitHub repository is publicly accessible at https://github.com/samlobo/FastHydroMap and have been deposited in Zenodo as archival snapshots.
